# Daily oscillation of the excitation/inhibition ratio is disrupted in two mouse models of autism

**DOI:** 10.1101/2021.07.05.451213

**Authors:** Michelle C.D. Bridi, Nancy Luo, Grace Kim, Caroline O’Ferrall, Ruchit Patel, Sarah Bertrand, Sujatha Kannan, Alfredo Kirkwood

## Abstract

Alterations to the balance between excitation and inhibition (E/I ratio) are postulated to underlie behavioral phenotypes in autism spectrum disorder (ASD) patients and mouse models. However, in wild type mice the E/I ratio is not constant, but instead oscillates across the 24h day. Therefore, we tested whether the E/I oscillation, rather than the overall E/I ratio, is disrupted in two ASD-related mouse lines: *Fmr1* KO and BTBR, models of syndromic and idiopathic ASD, respectively. The E/I ratio is dysregulated in both models, but in different ways: the oscillation is lost in *Fmr1* KO and reversed in BTBR mice. In both models these phenotypes associate with differences the timing of excitatory and inhibitory synaptic transmission and endocannabinoid signaling compared to wild type mice, but not with altered sleep. These findings raise the possibility that ASD-related phenotypes may be produced by a mismatch of E/I to the appropriate behavioral state, rather than alterations to overall E/I levels *per se*.

## INTRODUCTION

Autism spectrum disorder (ASD) is a prevalent neurodevelopmental disorder estimated to affect 1 in 54 children (Maenner et al., 2020). The core features of ASD are language deficits, social interaction deficits, and repetitive or restrictive behaviors and interests. These features are accompanied by sensory abnormalities in >90% of cases, and sensory processing differences have been reported to affect every sensory modality (reviewed in Geschwind, 2009; Robertson and Baron-Cohen, 2017). The visual system in particular is of great interest, as behavioral deficits have been tied to altered processing in the primary visual cortex (V1) of *Fmr1* knockout (KO) mice (Goel et al., 2018). Moreover, a recent analysis in humans concluded that regarding alterations in gene expression, “the primary visual cortex is the most affected region in ASD” (Berto et al., 2022).

ASD is genetically diverse, but elevation of the ratio between excitatory and inhibitory signaling (E/I ratio) in the brain has been proposed as a unifying mechanism (Rubenstein and Merzenich, 2003). Accordingly, higher E/I ratio and/or lower inhibition has been observed in sensory cortices of many ASD mouse models (Antoine et al., 2019; Banerjee et al., 2016; Bridi et al., 2017; Cellot and Cherubini, 2014; Chao et al., 2010; Gibson et al., 2008; Wallace et al., 2012; Zhang et al., 2014; Zhao and Yoshii, 2019). However, the opposite (decreased E/I ratio, increased inhibition, and/or decreased excitation) has also been reported (Dani et al., 2005; Delattre et al., 2013; Etherton et al., 2011; Harrington et al., 2016; Tabuchi et al., 2007; Unichenko et al., 2018).

One possible explanation for these opposing findings is that regulation of the E/I ratio, rather than the E/I ratio itself, is disrupted in ASD. In wild type (WT) mice, the E/I ratio in primary visual cortex (V1) changes over the course of the 24h light:dark cycle, such that it is low during the light (rest) phase (Bridi et al., 2020). Therefore, an apparent elevation of the E/I ratio in ASD models could either be due to an increased E/I at all times of day, or a dysregulated (e.g. flattened or phase-shifted) E/I oscillation. However, studies that measure the E/I ratio a single time of day cannot distinguish between these possibilities.

These considerations prompted us to examine possible E/I ratio dysregulation in two ASD-related mouse models with disparate genetic causes: the *Fmr1* knockout (*Fmr1* KO) mouse, which models Fragile X syndrome (The Dutch-Belgian Fragile X Consortium, 1994), the most frequent monogenic cause of intellectual disability and ASD in humans (Hagerman et al., 2017), and the BTBR *T+ Itpr3^tf^/J* (BTBR) mouse, an inbred line that models idiopathic ASD (McFarlane et al., 2008; Meyza and Blanchard, 2017). We show that the E/I ratio is dysregulated in both *Fmr1* KO and BTBR mice but in different ways: the oscillation is flattened in *Fmr1* KO mice and the timing of the oscillation is reversed in BTBR mice. Differences in endocannabinoid (eCB) signaling, but not sleep architecture, correspond to the patterns of E/I dysregulation in both lines, suggesting that eCB disruption could be a common feature across genetically diverse forms of ASD.

## RESULTS

### The E/I ratio is dysregulated in two mouse models of ASD

We first determined whether the E/I ratio is dysregulated across the 24-h light/dark cycle in *Fmr1* KO mice. We measured the E/I ratio by electrically stimulating in V1 layer 2/3 and recording excitatory and inhibitory synaptic responses in pyramidal cells lateral to the stimulating electrode (Figure 1A). We initially measured E/I at two time points: ZT0 and ZT12. As expected, in WT controls, the E/I ratio was higher at ZT0 (*t*_(51)_=3.92, *P*=0.0003, unpaired *t* test). In contrast, the E/I ratio in *Fmr1* KO animals was not different between ZT0 and ZT12 (*U*=312, *P*=0.22, Mann-Whitney U test), consistent with E/I dysregulation. To distinguish between a flattening vs altered timing of the E/I oscillation, we then included additional time points (ZT6, 18; Figure 1A). While the E/I ratio in *Fmr1* WT animals was higher when the animal had been in the dark phase (ZT0, 18), there was no time of day effect in *Fmr1* KO animals, indicating that the E/I oscillation is flattened (Figure 1B).

**Figure 1.**
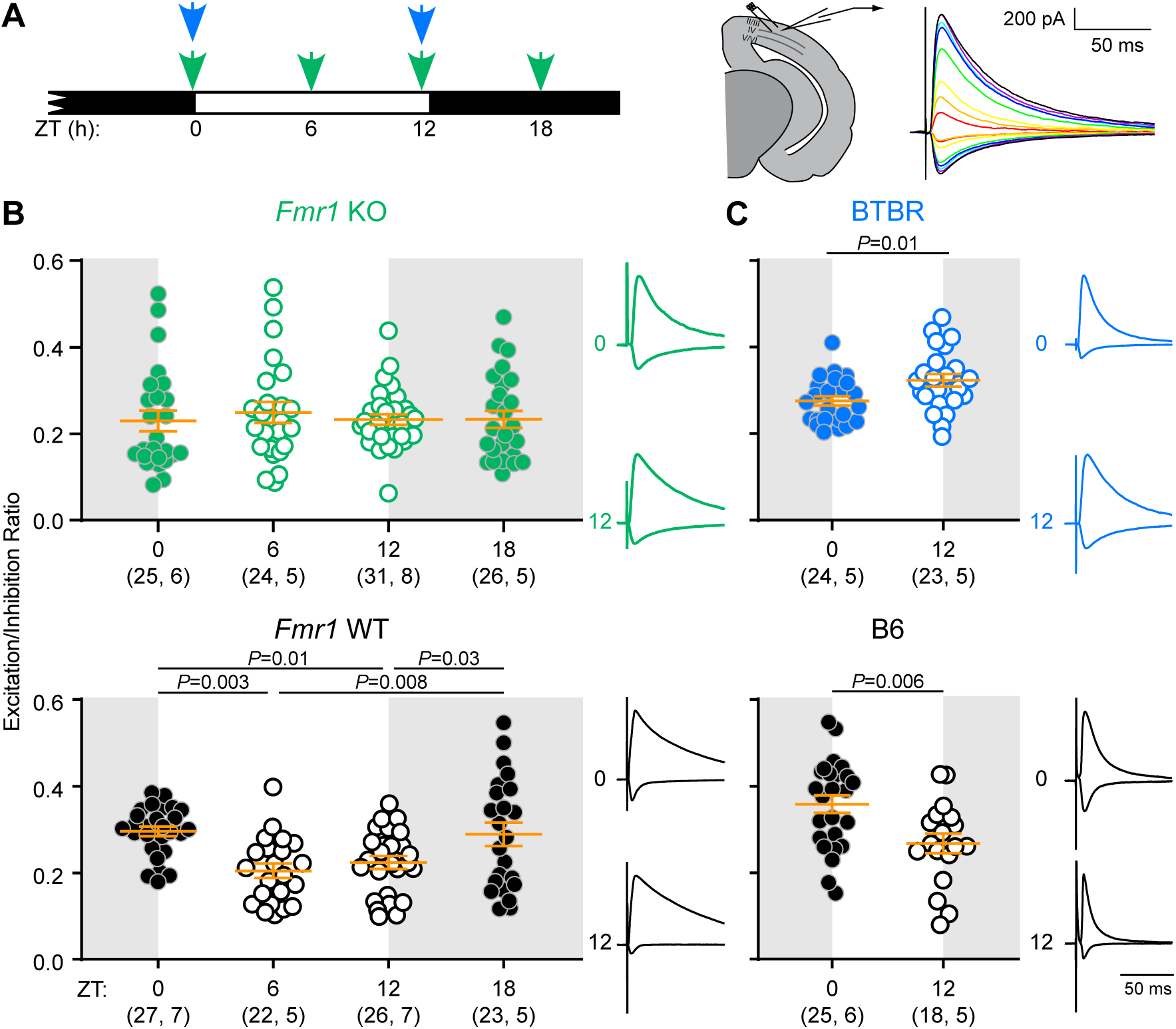
E/I is dysregulated in two mouse models of autism. (A) Left: *Fmr1* KO/WT (green) and BTBR/B6 (blue) mice were sacrificed at the indicated times of day. Right: Acute brain slices containing V1 were obtained, and responses to layer 2/3 stimulation over a range of intensities were recorded using whole-cell patch clamp of layer 2/3 pyramidal neurons. Inhibitory (upward deflection) and excitatory (downward deflection) responses were recorded in the same cell and the E/I ratio was calculated using the stimulation intensities over which the E/I ratio was stable (Bridi et al., 2020). (B) Oscillation of the E/I ratio across the day is absent in *Fmr1* KO mice. Top: *Fmr1* KO Kruskal-Wallis *P*=0.65. Bottom: *Fmr1* WT 1-way ANOVA Holm-Sidak post-hoc *P* values indicated. (C) BTBR mice exhibit E/I dysregulation, such that the E/I ratio is high at ZT12, contrasting with higher E/I at ZT0 in B6 mice. *P* values correspond to 2-tailed *t* tests. For all panels, sample size is indicated as (cells, mice). Error bars indicate mean±SEM. Example traces are normalized to peak inhibitory response.

To directly compare between genotypes, we conducted a 2-way ANOVA with time of day and genotype as main factors. We previously showed that the E/I ratio becomes stable within 4h of light/dark phase transitions, consistent with our current findings in *Fmr1* WT mice (Bridi et al., 2020); Figure 1). Therefore, for each genotype, data were pooled into dark (ZT0, 18) and light (ZT6, 12) phase groups. There was a significant interaction between genotype and time of day but no main genotype effect (Supplementary Table 1), confirming that E/I regulation, rather than overall magnitude, is altered in *Fmr1* KO mice. Post-hoc analysis revealed that the E/I ratio was significantly lower in the dark phase (*P*=0.003, Holm-Sidak test) but not significantly higher in the light phase (*P*=0.18) in *Fmr1* KO mice, indicating that the E/I oscillation flattening is predominantly carried by a decrease in the dark phase.

We then examined the E/I oscillation in BTBR and C57Bl/6J (B6) control mice (Figure 1A). As expected, B6 controls exhibited a higher E/I ratio at ZT0 than at ZT12 (Figure 1C). In contrast, the E/I ratio in the BTBR mice followed the opposite pattern, with a higher E/I ratio at ZT12 than ZT0 (Figure 1C). This indicates that the E/I ratio oscillates, but with reversed timing, in BTBR mice. A 2-way ANOVA revealed an interaction between genotype and time of day but no main effect of genotype (Supplementary Table 1), demonstrating that regulation of the E/I oscillation, rather than overall magnitude, is altered in BTBR mice. Post-hoc analysis showed that, compared to B6 controls, BTBR mice had a significantly lower E/I ratio at ZT0 (*P*=0.001, Holm-Sidak test) and higher E/I ratio at ZT12 (*P*=0.03), consistent with reversal of the E/I oscillation.

The E/I ratio oscillates over the 24h day in the lateral (layer 2/3-2/3) circuit, but not in the feedforward (layer 4-2/3) circuit in V1 of WT mice (Bridi et al., 2020). To confirm that *Fmr1* KO and BTBR mice did not display an abnormal oscillation in this circuit, we electrically stimulated layer 4 while recording synaptic currents in layer 2/3 of V1 slices. We confirmed that the E/I ratio did not change across the day for *Fmr1* KO, *Fmr1* WT, BTBR, or B6 animals (Supplementary Figure 1). In contrast with primary somatosensory cortex (S1) (Antoine et al., 2019), there was no main effect of genotype for either ASD model in the layer 4-2/3 circuit of V1 (Supplementary Figure 1; Supplementary Table 1). Additionally, V1 area is reduced in BTBR mice (Fenlon et al., 2015), raising the concern that V1 function may also be abnormal. Using optical imaging *in vivo,* we confirmed that V1, although smaller, is functional and expresses normal ocular dominance bias (Supplementary Figure 2). *Fmr1* KO mice are normal in these respects (Supplementary Figure 2).

### Altered sleep timing does not explain E/I dysregulation in two mouse models of ASD

Sleep regulates the oscillation of both excitatory and inhibitory synaptic transmission over the course of the day (Bridi et al., 2020; Liu et al., 2010). Therefore, it is possible that the observed E/I dysregulation in *Fmr1* KO and BTBR mice is due to abnormal sleep timing. Indeed, decreased total sleep and decreased numbers of rapid eye movement (REM) sleep bouts during the light phase have been reported in *Fmr1* KO mice (Boone et al., 2018; Saré et al., 2017); however, no studies to date have performed round-the-clock polysomnographic recordings in *Fmr1* KO or BTBR mice. We therefore continuously recorded EEG and EMG signals in the home cage using wireless telemetry for three days.

Sleep timing and architecture was grossly normal in *Fmr1* KO and BTBR mice compared to WT controls. Overall amounts of wake, non-REM (NREM) sleep, and REM sleep did not differ between *Fmr1* KO and WT or between BTBR and B6 mice (Supplementary Figure 3, Supplementary Table 2). We did not replicate the REM sleep deficit reported in *Fmr1* KO mice (Boone et al., 2018), possibly due to the longer recording sessions used in our study, or an emergence of the deficit with age (Saré et al., 2017). Furthermore, the distribution of time spent in each state across the day did not differ between *Fmr1* KO and WT mice (Figure 2A). The overall pattern of sleep timing appeared normal in BTBR mice, but there was a significant interaction between genotype and time of day for wake and REM amounts (Figure 2B, Supplementary Table 2). However, these slight differences cannot explain the reversal of the E/I oscillation in BTBR mice, since BTBR mice slept primarily during the light phase.

**Figure 2.**
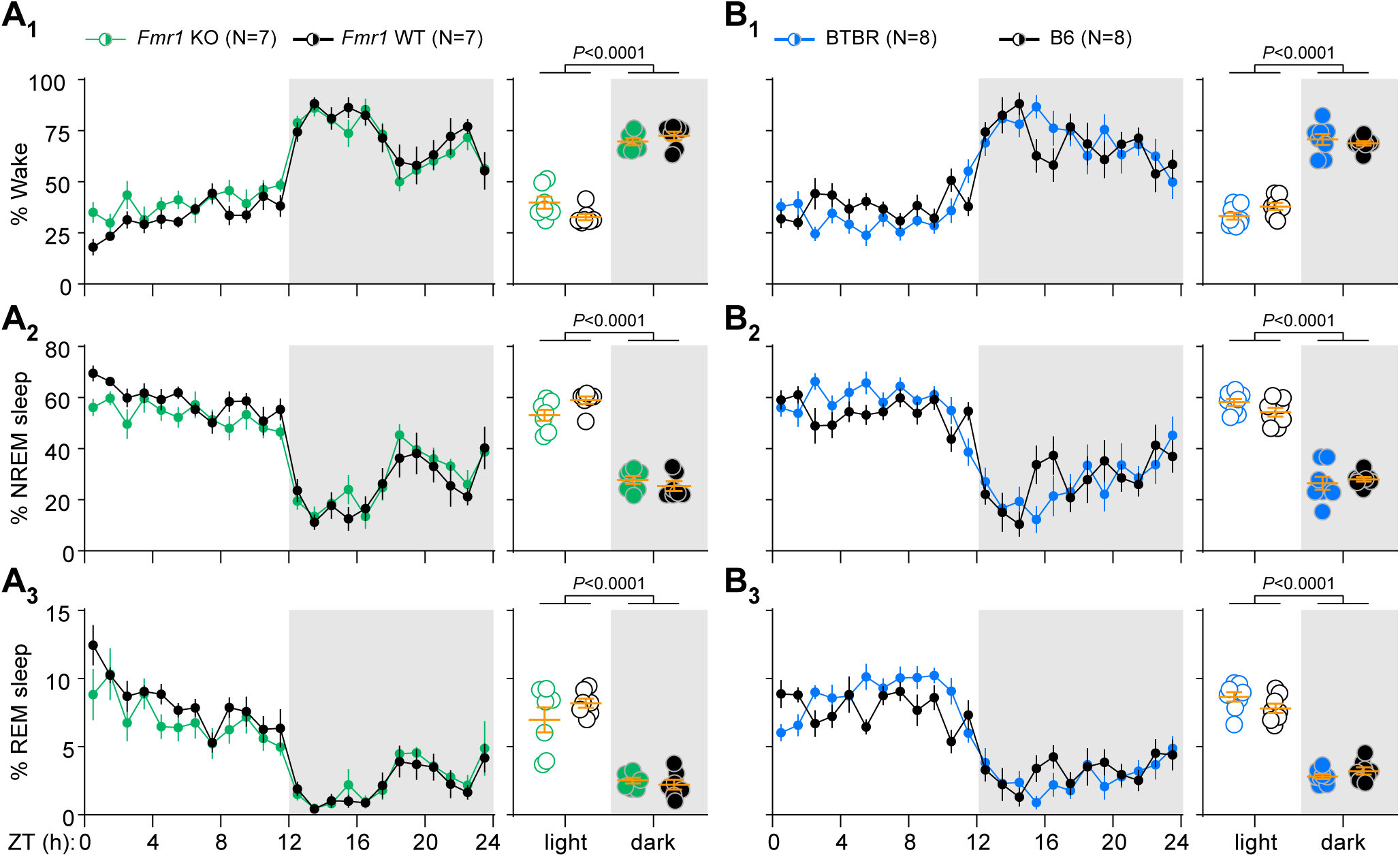
Sleep timing is normal in *Fmr1* KO and BTBR mice. (A) Percent time *Fmr1* mice spent awake (A_1_), in NREM sleep (A_2_), and in REM sleep (A_3_). There was no significant main effect of genotype or genotype × time of day interaction for any arousal state when data were broken into 1h bins (left) or averaged across the 12-h light and dark phases (right) (Supplementary Table 2). There was a significant main time of day effect for all states (Supplementary Table 2; \**P*<0.05 vs ZT0, Holm-Sidak post-hoc test). (B) Percent time BTBR and B6 mice spent in each arousal state. BTBR mice displayed grossly normal sleep timing (i.e. slept more during the light phase) and there was no significant main effect of genotype for any arousal state. There were small but significant interaction effects for wake (B_1_) and REM sleep (B_3_) when data were broken into 1h bins (left) and for REM sleep (B_3_) when data were averaged into 12h bins (right) (Supplementary Table 2). There was a significant main time of day effect for all states (Supplementary Table 2; *\*P*<0.05 vs ZT0, Holm-Sidak post-hoc test). Data were averaged over 3 recording days for each mouse prior to statistical analysis. All data were analyzed using 2-way repeated measures ANOVAs. Data are shown as mean ± SEM. N=number of mice.

We also examined the data for evidence of altered sleep quality in each line. Neither *Fmr1* KO nor BTBR mice had fragmented sleep (defined as shorter bout duration, more sleep bouts, and/or more sleep-wake transitions) (Supplementary Figure 3B-E). However, there were slight but significant alterations in spectral power during sleep. Most notably, NREM delta power was decreased in both models and REM theta power was increased in BTBR mice compared to B6 controls (Supplementary Figure 3E). We also observed a strong trend towards increased waking gamma power in *Fmr1* KO mice (Supplementary Figure 3F), consistent with previous reports (Goswami et al., 2019; Lovelace et al., 2018; Sinclair et al., 2017).

### Both excitation and inhibition are dysregulated in two mouse models of ASD

Our findings indicate that the E/I ratio is dysregulated across the day in two ASD-related mouse lines. However, these data do not reveal whether excitation, inhibition, or both are dysregulated. To determine this, we measured miniature excitatory and inhibitory postsynaptic currents (mEPSCs, mIPSCs) in V1 layer 2/3 at ZT0 and ZT12.

In *Fmr1* WT mice, mEPSC frequency was higher at ZT0 and mIPSC frequency was higher at ZT12 (Figure 3A, B; Supplementary Table 3). In contrast, mEPSC and mIPSC frequency in *Fmr1* KO mice was the same at both times of day (Figure 3A, B; Supplementary Table 3). 2-way ANOVAs with genotype and time of day as main factors (Supplementary Table 1) also revealed a significant main effect of genotype on mEPSC frequency, consistent with an increased density of excitatory synapses in *Fmr1* KO mice (Comery et al., 1997; Nimchinsky et al., 2001; The Dutch-Belgian Fragile X Consortium, 1994). No time-of-day differences in amplitude were observed for either mEPSCs or mIPSCs (Figure 3; Supplementary Table 3).

**Figure 3.**
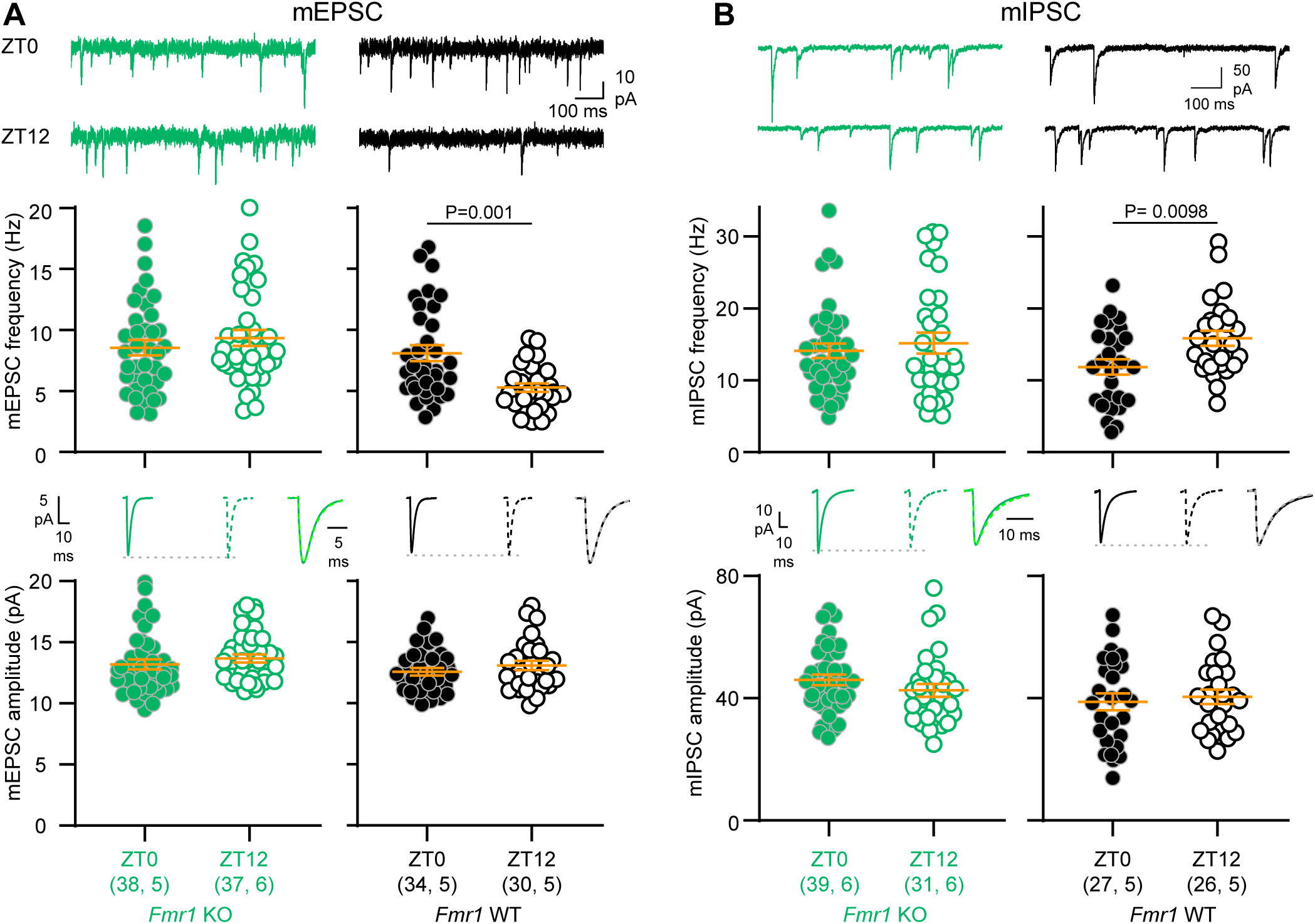
Oscillation of both excitation and inhibition is flattened in *Fmr1* KO mice. (A) mEPSC frequency does not change with time of day in *Fmr1* KO mice (*P*=0.35, Mann-Whitney test), but is higher at ZT0 in *Fmr1* WT control mice (*P*=0.001, Mann-Whitney test). Amplitude did not change with time of day in either genotype (KO: *P*=0.19; WT: *P*=0.39; Mann-Whitney test). (B) mIPSC frequency does not change with time of day in *Fmr1* KO mice (*U*=596, *P*=0.93, Mann-Whitney test), but is higher at ZT12 in *Fmr1* WT control mice (*t*_(51)_=2.69, *P*=0.0098, *t* test). Amplitude did not change with time of day in either genotype (KO: *t*_(68)_=1.26, *P*=0.21; WT: *t*_(51)_=0.46, *P*=0.65, *t* test). Data are presented as mean±SEM. Sample size is indicated as (cells, mice). Averaged traces: solid lines indicate ZT0, dotted lines indicate ZT12; scaled, superimposed averaged traces illustrate that there was no difference in kinetics between the two times of day (Supplementary Table 3).

We performed the same experiment for BTBR and B6 control mice. In BTBR mice, both mEPSC and mIPSC frequency followed a pattern opposite to B6 mice: mEPSC frequency was lower (Figure 4A) and mIPSC frequency was higher (Figure 4B) at ZT0 than at ZT12. 2-way ANOVAs showed significant genotype × time of day interaction effects for both mEPSCs and mIPSCs (Supplementary Table 1). There were no time-of-day differences in amplitude (Figure 4, Supplementary Table 3).

**Figure 4.**
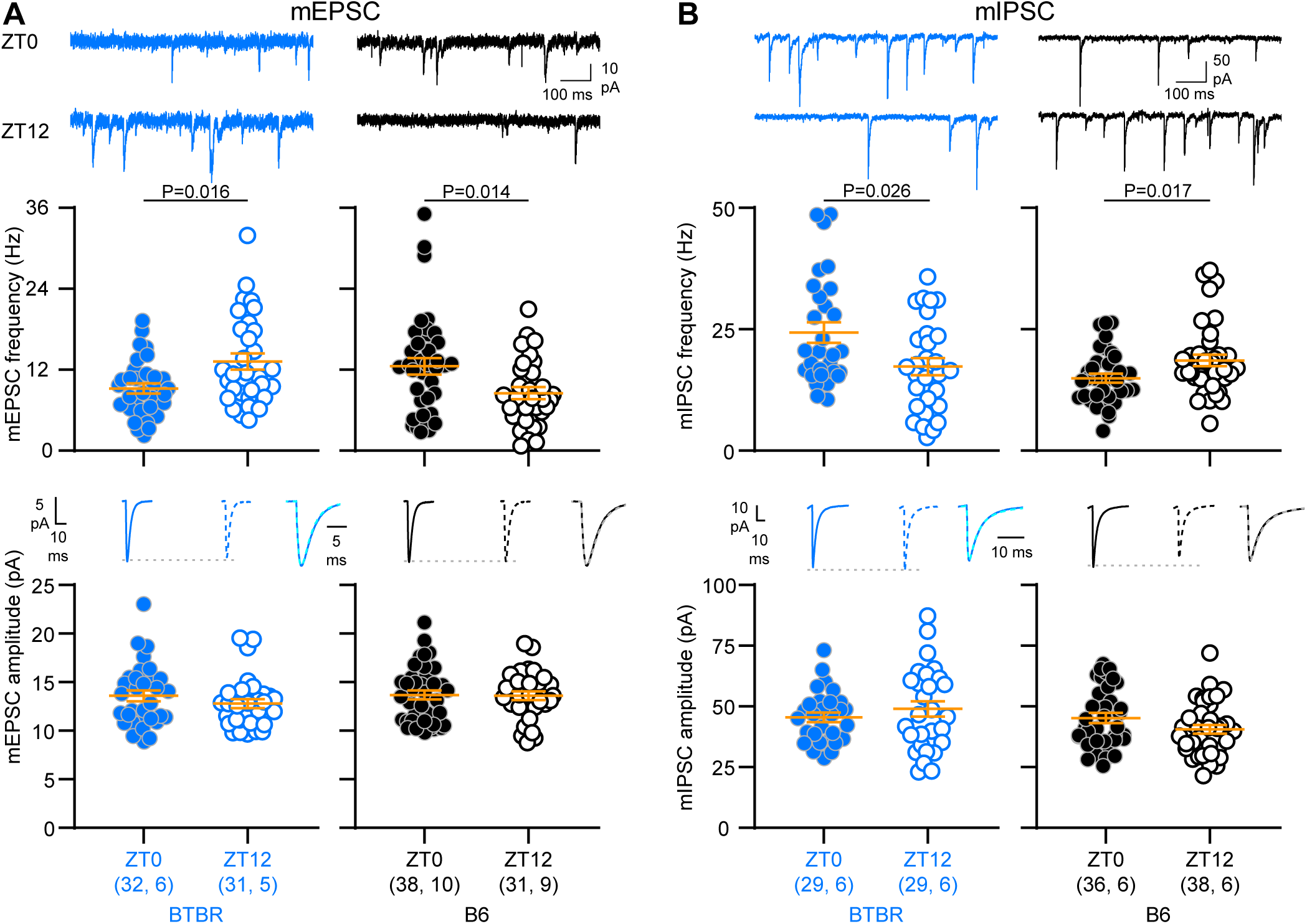
Oscillation of both excitation and inhibition is reversed in BTBR mice. (A) mEPSC frequency is higher at ZT12 in BTBR mice (*U*=322, *P*=0.016, Mann-Whitney test), and higher at ZT0 in B6 control mice (*t*_(67)_=2.53, *P*=0.014, *t* test). Amplitude did not change with time of day in either genotype (BTBR: *t*_(61)_=1.08, *P*=0.28; B6: *t*_(67)_=0.11, *P*=0.91; *t* test). (B) mIPSC frequency is higher at ZT0 in BTBR mice (*U*=278, *P*=0.026, Mann-Whitney test), and higher at ZT12 in B6 controls (*U*=464, *P*=0.017, Mann-Whitney test). Amplitude did not change with time of day in either genotype (BTBR: *t*_(56)_=0.93, *P*=0.36; B6: *t*_(72)_=1.71, *P*=0.09, *t* test). Data are presented as mean±SEM. Sample size is indicated as (cells, mice). Averaged traces: solid lines indicate ZT0, dotted lines indicate ZT12; scaled, superimposed averaged traces illustrate that there was no difference in kinetics between the two times of day (Supplementary Table 3).

mEPSC and mIPSC frequency track with the patterns of E/I dysregulation in each mouse line (Figure 1), indicating that both excitatory and inhibitory synaptic signaling underlie the dysregulation of the E/I ratio. Differences in mE/IPSC frequency, but not amplitude, across the day are in accordance with our previous study (Bridi et al., 2020) and are consistent with changes in synapse number and/or presynaptic machinery.

### eCB signaling is dysregulated in Fmr1 KO and BTBR mice

Multiple mechanisms can affect the E/I ratio (see Discussion). We previously identified eCB signaling as one mechanism contributing to time-of-day-dependent E/I changes in WT mice (Bridi et al., 2020). This raises the possibility that eCB signaling is altered in *Fmr1* KO and BTBR mice. In support of this idea, behavioral phenotypes in both lines can be normalized by pharmacological manipulation of eCB signaling (Wei et al., 2016). Therefore, we tested whether the time-of-day-specific sensitivity of inhibitory transmission to the eCB receptor agonist (+)-WIN 55,212-2 (WIN) is altered in each line. In WT mice, WIN suppresses spontaneous inhibitory currents (sIPSCs) when endogenous eCB levels are low (ZT12), and this effect is occluded when endogenous eCB levels are high (ZT0) (Bridi et al., 2020). In contrast, both *Fmr1* KO and BTBR mice showed altered patterns of WIN sensitivity. In *Fmr1* KO mice, WIN reduced sIPSCs at both times of day (Figure 5A). In BTBR mice, WIN suppressed sIPSCs when inhibition is abnormally high (ZT0) but this effect was occluded when inhibition was already low (ZT12) (Figure 5B). Repeating this experiment using an eCB receptor antagonist (SR141716A, SR) yielded complementary results (Supplementary Figure 4). Together, these results are consistent with flattening and reversal of inhibitory transmission in *Fmr1* KO and BTBR mice, respectively, that is driven in part by altered eCB signaling (Figure 6).

**Figure 5.**
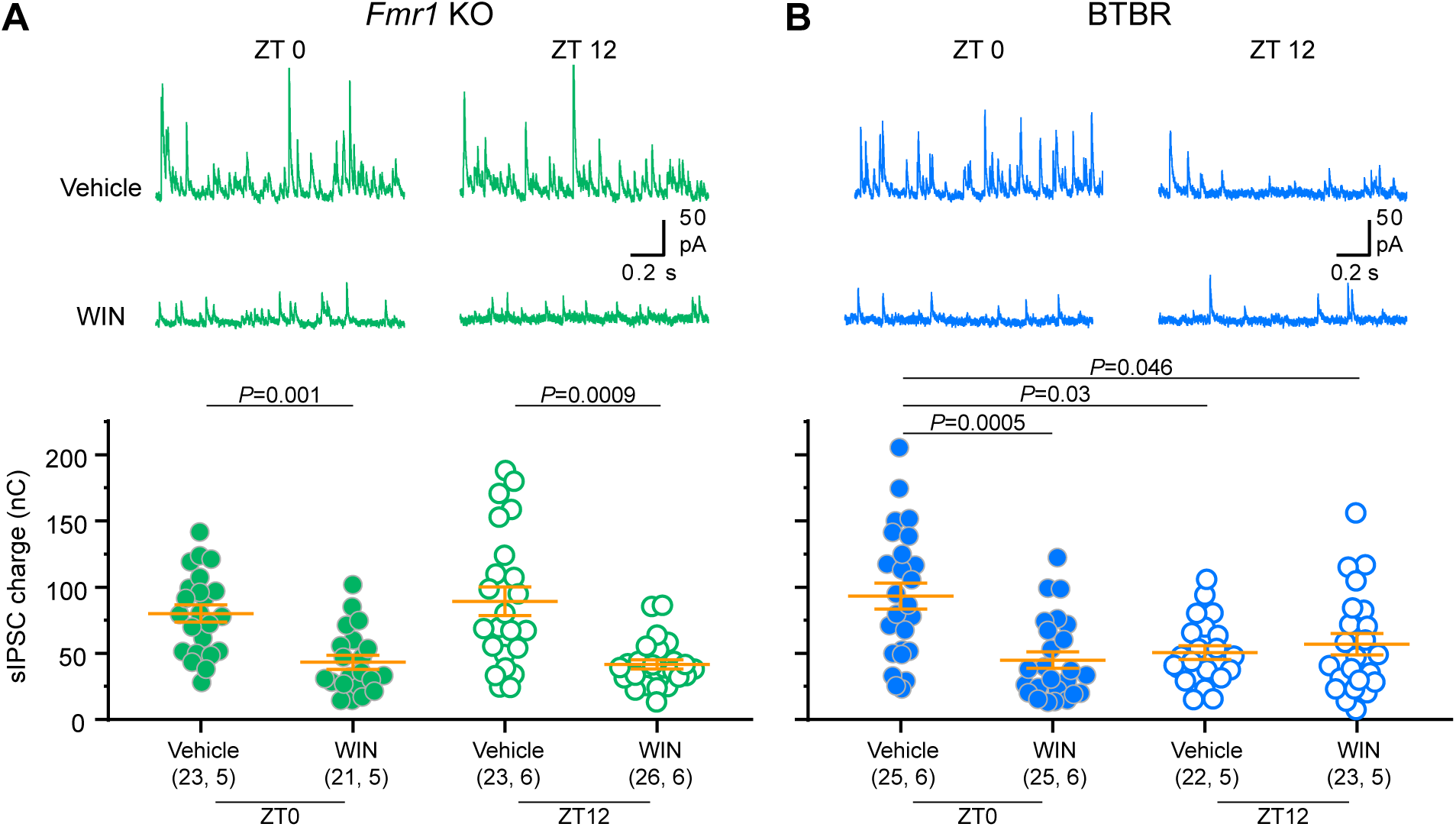
eCB signaling is dysregulated in *Fmr1* KO and BTBR mice. Slices were obtained at ZT0 or ZT12 and incubated in the eCB agonist WIN (10μM) or vehicle. sIPSCs were recorded from both treatment conditions in each animal. (A) Example traces (top) and quantification (bottom) of sIPSCs recorded from *Fmr1* KO slices. WIN significantly decreased sIPSC charge at both ZT0 and ZT12. sIPSC charge did not differ between times of day within vehicle or WIN treatment groups (P>0.9999). Kruskal-Wallis ANOVA on ranks *H*=28.73, *P*<0.0001. (B) Example traces (top) and quantification (bottom) of sIPSCs recorded from BTBR slices. WIN suppressed inhibitory transmission only at ZT0. Kruskal-Wallis ANOVA on ranks *H*=16.9, *P*=0.0007. Data are shown as mean ±SEM and sample size is indicated as (cells, mice). *P* values correspond to Dunn’s post-hoc test.

**Figure 6.**
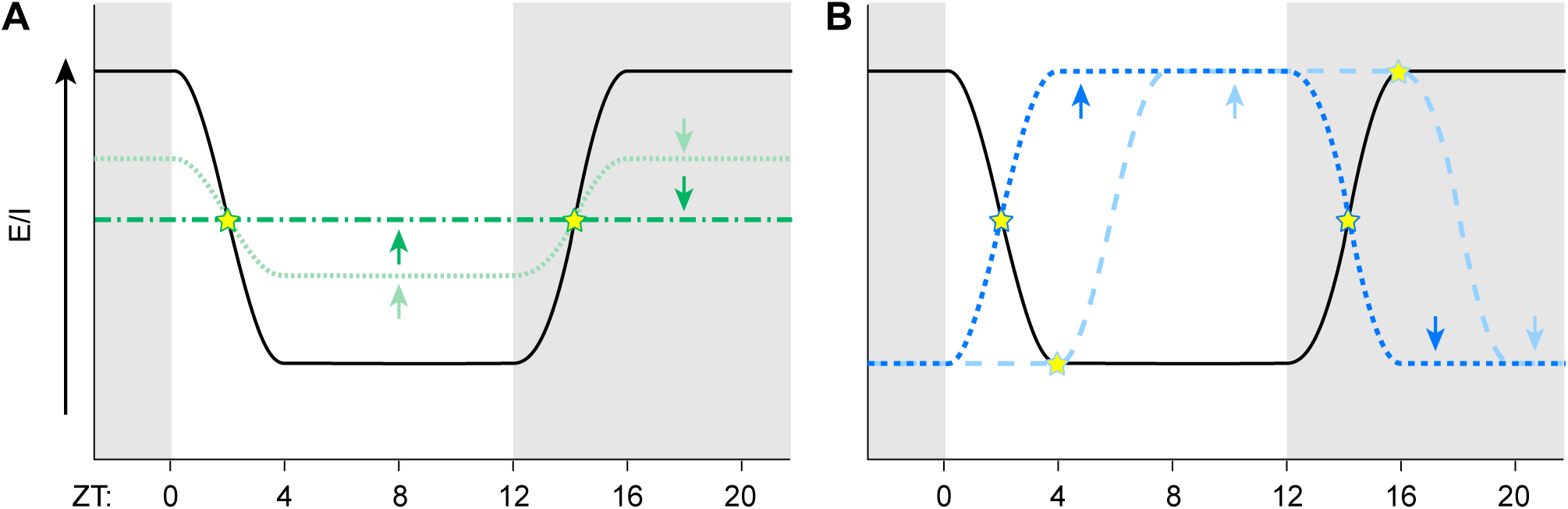
Altered timing or amplitude of the E/I oscillation preserves the WT E/I ratio at multiple times of day. (A) Compared to WT (black), decreased amplitude of the E/I oscillation (green) causes an apparent increase in E/I ratio during the light phase and/or decrease during the dark phase, but the E/I ratio is not different from WT at two times of day (stars). We observed amplitude flattening in *Fmr1* KO mice. (B) Compared to WT, altered timing such as a phase shift of the E/I oscillation can cause an apparent increase in the E/I ratio during the light phase, but the E/I ratio is not different from WT at multiple times of day (stars). We observed reversed timing of the oscillation in BTBR mice.

## DISCUSSION

In WT mice, the E/I ratio and correlations between ionic conductances vary over the 24h day, indicating daily rearrangement of connectivity and function under normal conditions (Bridi et al., 2020; Tran et al., 2019; Zong et al., 2023). Therefore, it is crucial to consider neuronal dysfunction in models of neurological conditions against a backdrop of dynamic circuit properties. In this study, we show that the daily E/I oscillation is dysregulated due to changes in both excitatory and inhibitory synaptic transmission in two mouse lines associated with ASD. The oscillation is flattened in *Fmr1* KO, and reversed, consistent with altered phase and/or periodicity, in BTBR mice. These changes cannot be explained by sleep timing, but patterns of inhibition are consistent with altered eCB signaling. Exploring how this dysregulation occurs is an important avenue for future investigation.

### Implications for ASD

A prominent hypothesis in the field states that elevated E/I ratio is a common mechanism leading to the behavioral phenotypes across genetically diverse forms of ASD (Rubenstein and Merzenich, 2003). We propose to refine this viewpoint: on average, the magnitude of the E/I ratio is normal, but it is elevated/lowered at inappropriate times of day (Figure 6). This may result in a mismatch between E/I and brain state that affects behavioral phenotypes.

Identifying the patterns of E/I dysregulation specific to each ASD-related mouse line may reconcile discrepancies across studies. The E/I ratio is low during the light (rest) phase in WT animals (Bridi et al., 2020). Therefore, measuring the E/I ratio only during the light phase predisposes studies to report an elevated E/I ratio, when there may be no overall difference, in mouse lines associated with ASD (Figure 6). Consistent with dysregulation, lower mIPSC frequency has been reported in V1 L2/3 of *Cntnap2*-, *Tsc1-,* and *Ube3a-*deficient mice (Bridi et al., 2017; Wallace et al., 2012; Zhao and Yoshii, 2019). In contrast, *Fmr1* KO mice have normal mIPSC frequency and spontaneous activity in V1 L2/3 (Goel et al., 2018; Zhong et al., 2018), which may be attributable to differences in the time of day at which recordings were conducted and/or genetic background.

In this study, we examined V1 because the E/I ratio oscillation in WT mice is best characterized in this region. However, the E/I ratio also undergoes a daily oscillation in hippocampus and medial prefrontal cortex (Bridi et al., 2020), raising the possibility that E/I dysregulation may extend to other brain regions. Consistent with this idea, the E/I ratio is elevated in CA1 of *Fmr1* KO and BTBR mice (Cellot et al., 2016; Han et al., 2014; Sabanov et al., 2017; Wahlstrom-Helgren and Klyachko, 2015), but whether these findings are due to dysregulation remains to be seen. Regardless, our positive results in V1 confirmed our central hypothesis of E/I dysregulation.

Furthermore, the E/I ratio oscillation is specific to the layer 2/3-2/3, but not the layer 4-2/3, pathway in WT mice (Supplementary Figure 1; Bridi et al., 2020). Here, we show that the E/I ratio remains fixed in the layer 4-2/3 circuit of both *Fmr1* KO and BTBR mice. Notably, the E/I ratio was not elevated in this circuit in either line, suggesting that pathways in which the E/I ratio is dynamically regulated are more likely to be affected in ASD models. However, the E/I ratio is elevated in S1 layer 4-2/3 of *Fmr1* KO mice (Antoine et al., 2019), raising the possibility that the circuit-specificity of the E/I ratio oscillation may differ across cortical regions.

### Candidate Mechanisms of E/I dysregulation

One intriguing implication of our findings is that regulation of excitatory and inhibitory synaptic transmission go hand-in-hand, since they are jointly dysregulated in both *Fmr1* KO and BTBR mice. This may occur via a central mechanism controlling both excitation and inhibition, or by the dependence of excitation on inhibition (or vice versa). While the exact mechanisms remain to be elucidated, changes in mE/IPSC frequency are consistent with formation/elimination of synapses across the 24h day, implicating a number of potential candidate mechanisms.

Four moving parts comprise the E/I ratio oscillation in WT mice: increased excitation/decreased inhibition during the active phase, and decreased excitation/increased inhibition during the rest phase. During the active phase, long-term potentiation has been implicated in increased excitatory transmission, while eCB signaling, visual experience, and cholinergic signaling control decreased inhibitory transmission. During the rest phase, sleep is required for decreased excitation and increased inhibition (Bridi et al., 2020; Liu et al., 2010; Vyazovskiy et al., 2008; Zong et al., 2023). The ways in which these mechanisms operate in WT mice remain poorly understood, limiting our ability to interpret the exact mechanisms of dysregulation in *Fmr1* KO and BTBR mice. Nevertheless, we examined how two of these mechanisms, sleep and eCB signaling, relate to E/I dysregulation.

Sleep disruption is a common feature of ASD (reviewed in Schreck and Richdale, 2020). However, despite a variety of altered sleep and activity characteristics, a grossly normal nocturnal activity pattern in preserved across many commonly used ASD-related mouse lines (Angelakos et al., 2017; Colas et al., 2005; Ehlen et al., 2015; El Helou et al., 2013; Ingiosi et al., 2019; Li et al., 2015; Lipton et al., 2017; Liu et al., 2017; Lu et al., 2019; Saré et al., 2021, 2017; Seok et al., 2018; Thomas et al., 2017; Tsuchiya et al., 2015; Wither et al., 2012; Zhang et al., 2008). Here we confirmed that sleep timing follows a normal pattern of nocturnal activity in *Fmr1* KO and BTBR mice. Therefore, dysregulation is unlikely to arise from altered sleep timing. However, NREM delta power was reduced in both genotypes, consistent with reports in other mouse lines (Angelakos et al., 2017; Ehlen et al., 2015; Ingiosi et al., 2019; Johnston et al., 2014; Liu et al., 2017; Lu et al., 2019), suggesting that altered neuronal activity during NREM sleep could contribute to E/I dysregulation.

eCB signaling has also been implicated in ASD in humans (Aran et al., 2019; Pretzsch et al., 2019), as well as in *Fmr1* KO, BTBR, and other ASD-related mouse lines (Busquets-Garcia et al., 2013; Földy et al., 2013; Gomis-González et al., 2016; Jung et al., 2012; Maccarrone et al., 2010; Qin et al., 2015; Speed et al., 2015; Wei et al., 2016; Zhang and Alger, 2010). Here, we report that inhibitory transmission is susceptible to manipulations of eCB signaling in *Fmr1* KO and BTBR mice, indicating that eCB signaling pathways are operational. However, the response to eCB manipulation is flattened in *Fmr1* KO and reversed in BTBR mice, consistent with altered timing of eCB release and/or eCB receptor expression contributing to dysregulation of inhibition in both lines.

An additional candidate mechanism is glial function. Microglia and astrocytes play a role in pruning and function of excitatory and inhibitory synapses (Blinzinger and Kreutzberg, 1968; Chung et al., 2013; Favuzzi et al., 2021; Lee et al., 2021; Oliet et al., 2001; Paolicelli et al., 2011; Shigetomi et al., 2012), thus shaping connectivity in the cortex (Liu et al., 2021). Importantly, microglial and astrocyte function, which in WT mice depend on arousal state (Bellesi et al., 2017; Ding et al., 2013; Ingiosi et al., 2020; Liu et al., 2019; Paukert et al., 2014; Stowell et al., 2019), is altered in both *Fmr1* KO and mice BTBR mice (Eissa et al., 2020; van der Goot et al., 2019; Heo et al., 2011; Higashimori et al., 2013; Jawaid et al., 2018). Whether glia play a role in E/I dysregulation, however, remains to be explored.

### Consequences of dysregulated E/I balance

The exact behavioral impact of altered E/I balance in ASD remains largely unclear. A common view states that network hyperexcitability associated with reduced inhibition and altered inhibitory plasticity impairs neural processing (for recent reviews see Antoine, 2022; Chen et al., 2022; Ferguson and Gao, 2018; Liu et al., 2022), an idea well suited to account for perceptual learning deficits resulting from impairments in sensory discrimination (Goel et al., 2018). Alternatively, altered excitatory and inhibitory synaptic transmission may not be a primary effect of the ASD phenotype, but rather reflect homeostatic compensations that mitigate other types of dysfunction (Antoine et al., 2019; Domanski et al., 2019; Nelson and Valakh, 2015). E/I dysregulation might offer unique opportunities to discriminate between these two possibilities because E/I differences between mutant and WT animals will vary in magnitude and sign across the day. In these cases, time-of-day dependent variations in the degree of behavioral impairments, as reported recently (Sawicka et al., 2019), correlating with daily E/I discrepancies will support a direct impact of altered E/I on behavioral phenotype.

### Conclusions

The E/I ratio is dynamically regulated, likely via multiple mechanisms acting in concert. Each of these mechanisms has the potential to be affected by a variety of genetic and environmental factors, making the E/I oscillation vulnerable to disruption. Therefore, E/I dysregulation may be a point of convergence for genetically diverse forms of ASD, ultimately resulting in common behavioral phenotypes. Further exploring the underlying mechanisms will provide crucial insight to the pathophysiology of ASD.

## METHODS

### Animal

All procedures were approved by the Johns Hopkins University Institutional Animal Care and Use Committee. Mice were bred in-house on a 12:12 light:dark cycle (lights on at 7AM). Upon weaning, naïve mice were entrained for at least two weeks to a 12:12 light:dark cycle in custom entrainment chambers, with the timing of lights-on adjusted according to the experiment. Due to X-linkage of the *Fmr1* gene, only male *Fmr1* WT and KO mice were used. C57Bl/6J and BTBR mice of either sex were used. Littermates were distributed across experimental groups.

### Slice Preparation

300 μm thick coronal brain slices containing V1 were prepared as described previously (Bridi et al., 2020). Briefly, slices were cut in ice-cold dissection buffer containing 212.7 mM sucrose, 5 mM KCl, 1.25 mM NaH_2_PO_4_, 10 mM MgCl_2_, 0.5 mM CaCl_2_, 26 mM NaHCO_3_, and 10 mM dextrose, bubbled with 95% O_2_/5% CO_2_ (pH 7.4). Slices were transferred to normal artificial cerebrospinal fluid (similar to the dissection buffer except that sucrose was replaced by 119 mM NaCl, MgCl_2_ was lowered to 1 mM, and CaCl_2_ was raised to 2 mM) and incubated at 30°C for 30 min and then at room temperature for at least 30 min before recording.

### Whole-Cell Recording

Visualized whole-cell voltage clamp recordings were made from pyramidal neurons in L2/3 (35% depth from the pia) of V1. In slices from BTBR mice, recordings were restricted to a smaller posterior/medial region to account for the size and location of V1 in these mice (Fenlon et al., 2015; Supplementary Figure 2). Glass pipette recording electrodes (3–6 MΩ) were filled with different internal solutions according to each experiment, all of which were adjusted to pH 7.2– 7.3, 280–295 mOsm. Cells with an input resistance ≥ 150 MΩ and access resistance ≤ 25 MΩ were recorded. For all whole cell recordings, cells were discarded if these values changed more than 25% during the experiment. Data were filtered at 2 kHz and digitized at 10 kHz using Igor Pro (WaveMetrics, Portland, OR).

#### E/I ratio

The internal pipette solution for recording evoked EPSCs and IPSCs contained 8mM KCl, 125 mM cesium gluconate, 10 mM HEPES, 1 mM EGTA, 4mM Mg-ATP, 0.5mM Na-GTP, and 5 mM QX-314. Responses were recorded in the presence of 100 μM DL-APV. Reversal potentials for excitatory and inhibitory currents of +10 mV and -55 mV (without junction potential compensation) were used (Bridi et al., 2020). To evoke synaptic responses, a double-barrel glass stimulating pipette filled with ACSF was placed approximately 100-200 μm lateral to the recording electrode (layer 2/3 stimulation) or in the middle of the cortical thickness (layer 4 stimulation). A series of stimulations over a range of intensities was delivered, and the responses over the range of intensities producing a stable E/I ratio were used (Bridi et al., 2020).

#### Miniature postsynaptic current recordings

For mEPSC recordings, 1 μM TTX, 100 μM DL-APV and 2.5 μM gabazine were added to the perfusion buffer to isolate AMPAR-mediated mEPSCs. An internal pipette solution containing the following ingredients was used: 8 mM KCl, 125 mM cesium gluconate, 10 mM HEPES, 1 mM EGTA, 4 mM NaATP, and 5 mM QX-314. To record mIPSCs, 1 μM TTX, 100 μM DL-APV and 20 μM CNQX were included in the bath. The internal pipette solution contained: 8 mM NaCl, 120 mM cesium chloride, 10 mM HEPES, 2 mM EGTA, 4 mM MgATP, 0.5 mM NaGTP, and 10 mM QX-314. V_m_ was held at -70 mV.

#### Spontaneous IPSC recordings

To record sIPSCs, conditions were similar to mEPSC recording except no QX-314 was included in the internal pipette solution, V_m_ was held at +10 mV, and the bath contained only 10μM (+)-WIN55,212-2 (Cayman Chemical, Ann Arbor, MI), 10μM SR 141716A (Abmole, Houston, TX), or vehicle (0.1% DMSO). Slices were pre-incubated in these drugs for at least 1h prior to recording. Control and drug-treated slices were obtained from the same animals.

### Polysomnography recording

#### Surgery

4-5 week old mice were placed under isoflurane (1-2%) anesthesia and immobilized. A pocket was formed under the skin and a wireless transponder attached to four recording leads (model HD-X02, Data Sciences International, St. Paul, MN) was inserted into the pocket. Two EMG leads with 0.5 cm exposed wire were inserted into the cervical trapezius muscles and held in place with 5-0 silk sutures. The skull was cleaned with H_2_O_2_ and two holes were drilled for EEG lead placement (location posterior to bregma/lateral to the midline: 3.4mm/2.5mm and 1mm/1mm). EEG leads with 1-2 mm exposed wire were inserted into the holes to make contact with the dura and affixed with dental acrylic and cyanoacrylate glue. The wound was sutured closed and treated with triple antibiotic ointment. Mice were allowed to recover in their home cage for at least 7 days before recording. Home cages were then placed on top of telemetry receiver pads and EEG and EMG signals were captured (Data Sciences International, PhysioTel Receiver RPC-1) using Ponemah software (Data Sciences International, Version 6.40) for 3 consecutive days on a light-dark cycle matching the vivarium environment. The EEG/EMG signal was sampled at 500 Hz.

### Optical imaging of the intrinsic cortical signal

Animals were anesthetized using isoflurane in O_2_ (induction: 2-3%, maintenance: 0.5-1%) supplemented with chlorprothixene (2 mg/kg i.p.). An incision was made in the scalp and lidocaine was applied to the margins. Exposed skull above V1 was covered in 3% agarose and an 8mm round glass coverslip. Surface vasculature was visualized by illuminating the area with 555nm light. Then the camera was focused 600 μm below the cortical surface and the area was illuminated with 610 nm light. A high refresh rate monitor (1024 × 768 @120 Hz; ViewSonic, Brea, CA) was aligned in the center of the mouse’s visual field 25cm in front of the eyes. Visual stimuli consisted of a white horizontal bar on a black background (2° height) presented either to the entire visual field or only to the binocular visual field (-5° to +15° azimuth), moving continuously upward or downward (5 minutes per direction). Cortical signals were imaged using a Dalsa 1M30 CCD camera (Dalsa, Waterloo, Canada).

### Quantification and statistics

#### Whole-cell recordings

##### E/I ratio

Peak response amplitude at each holding potential was measured for each stimulus intensity and the ratio between the excitatory and inhibitory peak was calculated (Igor Pro). If multiple peaks were observed in the postsynaptic response, the magnitude of the first peak was used in order to limit the analysis to monosynaptic responses. If the first peak could not be clearly resolved, the cell was discarded from the analysis.

##### Spontaneous synaptic events

mEPSCs and mIPSCs were analyzed using the MiniAnalysis program (Synaptosoft, Decatur, GA). Only cells with root mean square (RMS) noise <2 (mEPSCs) or <4 (mIPSCs) were included in the analysis and event detection threshold was set at 3 times the RMS noise. 300 events with rise time <3 msec (mEPSCs) or <5 msec (mIPSCs) were selected for each cell to calculate frequency and amplitude. Non-overlapping events were used to construct the averaged traces. Spontaneous IPSCs were analyzed by calculating the unit charge (nA/s) with custom code (Matlab, MathWorks, Inc, Natick, MA) (Bridi et al., 2020). The baseline was calculated and subtracted for each 500 msec of recording. Charge was calculated as the integral of the baseline-subtracted signal. 3-4 minutes of recording were quantified for each cell.

#### Polysomnography

Arousal stages were scored manually off-line as NREM sleep, REM sleep, and wake by a trained experimenter (MB) in 4-s epochs (SleepSign for Animal, Kissei Comtec). The percent time spent in each state, along with number and duration of bouts, was calculated. Power spectra were computed within each arousal state by performing a fast Fourier transform on the EEG signal with 0.5 Hz resolution. For each arousal state, spectra were normalized to the total EEG power (0.5-80 Hz) in that state. One B6 and one BTBR animal were included in sleep architecture analysis but excluded from spectral analysis due to differences in electrode placement.

#### Optical imaging of the intrinsic cortical signal

The cortical response at the stimulus frequency was extracted by Fourier analysis. Images were smoothed by a 5×5 low-pass Gaussian filter and the binocular region of interest (ROI) was defined as the 70% of pixels with the highest intensity in the ipsilateral eye map (Matlab, Mathworks, Natick, MA). The average number of pixels activated by the full visual field stimulus was calculated as a relative measure of V1 size. The ocular dominance value of each pixel in the binocular region was calculated as (contra-ipsi)/(contra+ipsi) and averaged to obtain the ODI.

#### Statistics

Data were analyzed with 2-tailed unpaired *t*-tests, Mann-Whitney tests, 2-way ANOVAs (±repeated measures) with Holm-Sidak posthoc analysis, or Kruskal-Wallis with Dunn’s posthoc analysis, as indicated in the figure and table legends (GraphPad Prism, San Diego, CA). *P<*0.05 was considered significant. In cases where data were not normally distributed (Kolmogorov-Smirnov normality test), nonparametric tests were used. Sample size is displayed in the figures as (number of cells, number of animals). Lines and error bars in all figure dot plots indicate mean and SEM.

## Supplementary Figure Legends

**Supplementary Figure 1.**
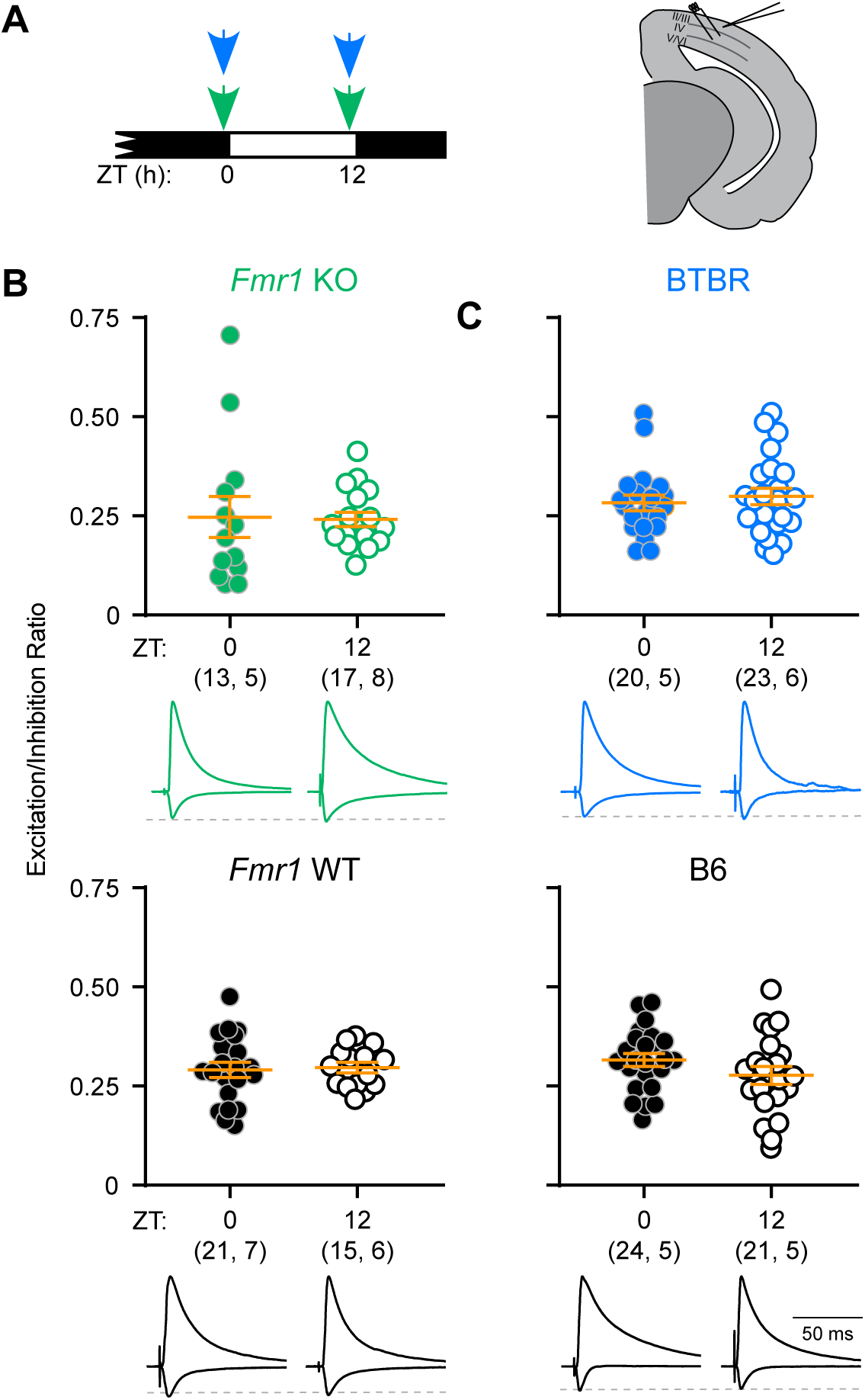
The layer 4-2/3 E/I ratio is not affected by time of day. (A) *Fmr1* WT, *Fmr1* KO, B6, and BTBR mice were sacrificed at two times of day and acute brain slices containing V1 were collected for whole-cell patch clamp recordings of layer 2/3 pyramidal neurons in response to layer 4 stimulation. (B, C) The E/I ratio was not different between ZT0 and ZT12 in *Fmr1* KO, BTBR, or WT mice (2-tailed *t* tests within genotype; Supplementary Table 1). When each line was compared to its WT control using 2-way ANOVAs, no significant main effects of time, genotype, or time × genotype interaction were observed (Supplementary Table 1). Sample size is indicated as (cells, mice). Error bars indicate mean±SEM. Example traces show the inhibitory response (upward deflection) and excitatory response (downward deflection) in the same cell and are normalized to peak inhibitory response.

**Supplementary Figure 2.**
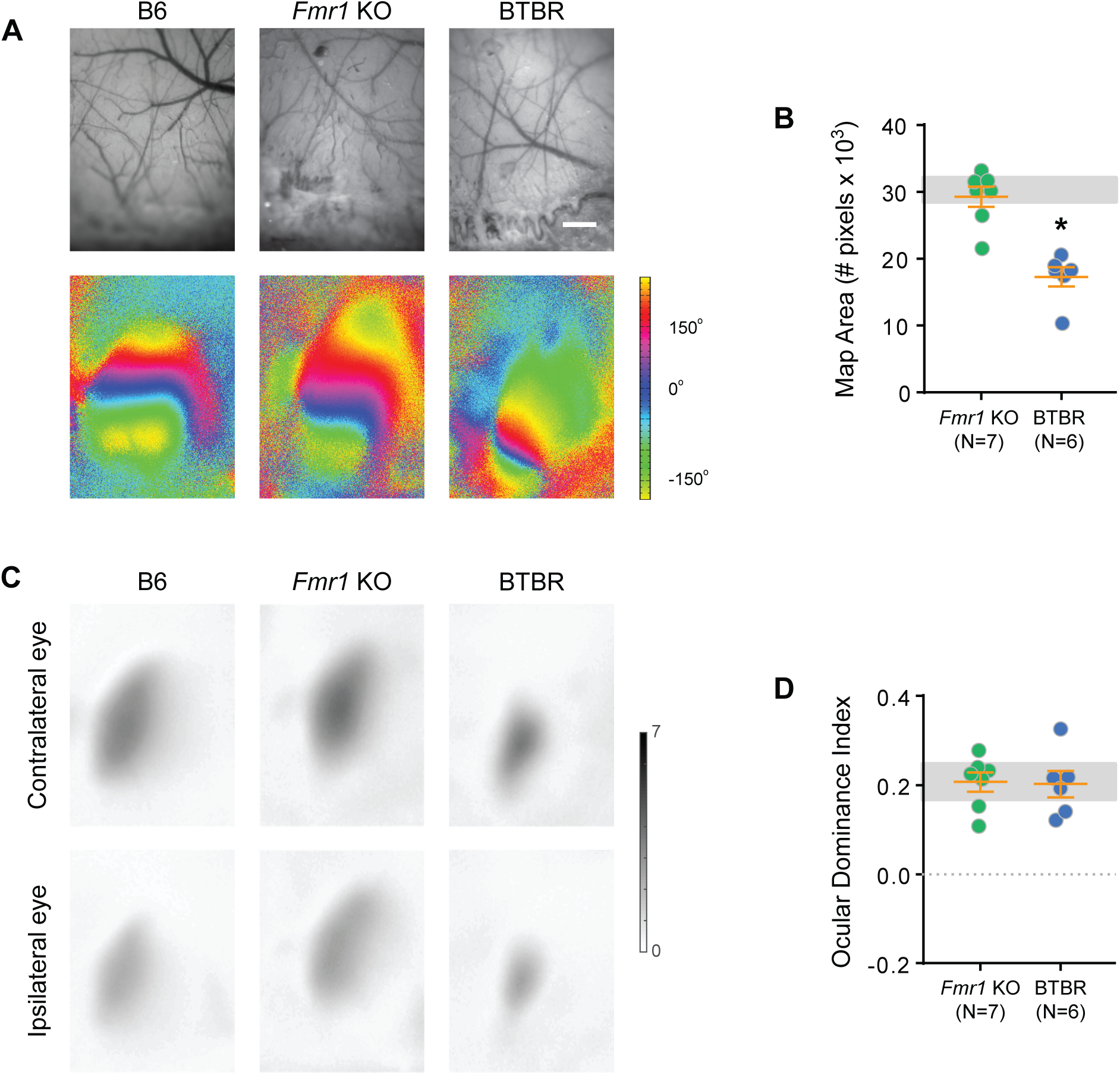
V1 is functional and expresses normal ocular dominance bias in *Fmr1* KO and BTBR mice. **(A)** Example images of the cortical surface vasculature (top) and retinotopic maps (bottom) obtained by optical imaging of the intrinsic cortical signal while presenting a visual stimulus to the entire visual field of both eyes simultaneously. V1 of both mouse lines was functional and displayed retinotopic organization. Scale bar: 1mm. **(B)** V1 size was normal in *Fmr1* KO mice but smaller in BTBR mice, compared with WT controls (shaded gray region indicates mean ± 95% CI of B6 mice). Kruskal-Wallis ANOVA on ranks *P*=0.0011; *Fmr1* KO *P*>0.999, BTBR *P*=0.0007, Dunn’s post-hoc test vs. B6). **(C)** Example images showing the magnitude of response to each eye during presentation of a visual stimulus to the binocular visual field. **(D)** The ocular dominance index in binocular V1 was normal in both *Fmr1* KO and BTBR mice, compared to WT controls (shaded gray region indicates mean ± 95% CI of B6 mice). ANOVA *F*_(2, 25)_=0.014, *P*=0.986. Sample size is indicated as (# mice) and error bars represent SEM.

**Supplementary Figure 3.**
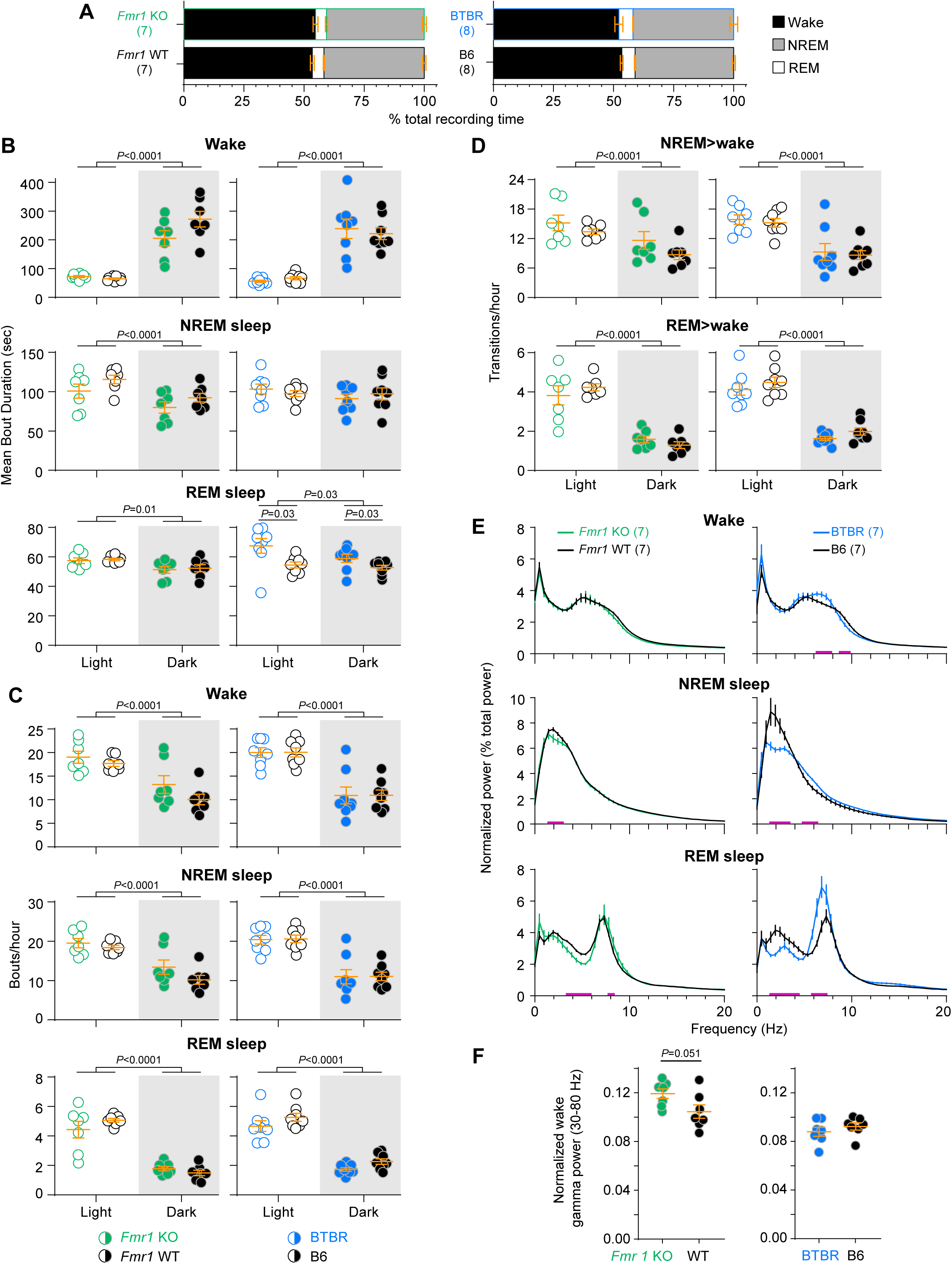
Sleep architecture and EEG power in ASD model mice. **(A)** Overall amounts of wake, NREM sleep, and REM sleep did not differ between *Fmr1* KO and WT mice, or between BTBR and B6 mice (2-tailed *t* test *P>*0.05; Supplementary Table 2). **(B)** Sleep/wake bout durations did not differ between *Fmr1* KO and WT mice. Both *Fmr1* KO and WT mice had shorter wake and longer sleep bouts during the light phase. BTBR mice, on the other hand, showed subtle changes in sleep architecture. REM sleep bouts were significantly longer in both the light and dark phases compared to B6. Both genotypes had shorter wake and longer REM bouts during the light phase. *P* values indicate significance on 2-way repeated measures ANOVAs; Supplementary Table 2. **(C,D)** All arousal states were more fragmented (more bouts and sleep-wake transitions) during the light phase than during than during the dark phase. No main effects of genotype or genotype × time interactions were observed. *P* values indicate significance on 2-way repeated measures ANOVAs (Supplementary Table 2). **(E)** Power spectra were calculated separately for wake, NREM sleep, and REM sleep arousal states and normalized to total spectral power (0.5-80 Hz) within that state. Data were compared using 2-way ANOVAs with genotype and frequency as main factors (Supplementary Table 2). Magenta lines indicate the frequencies at which genotypes are significantly different (Holm-Sidak post-hoc test *P<*0.05). **(F)** Wake power in the gamma range showed a strong trend toward higher power in *Fmr1* KO than WT controls, but no difference between B6 and BTBR mice (2-tailed *t* test; Supplementary Table 2). For all panels, sample size is indicated as (mice) and bars represent mean ± SEM.

**Supplementary Figure 4.**
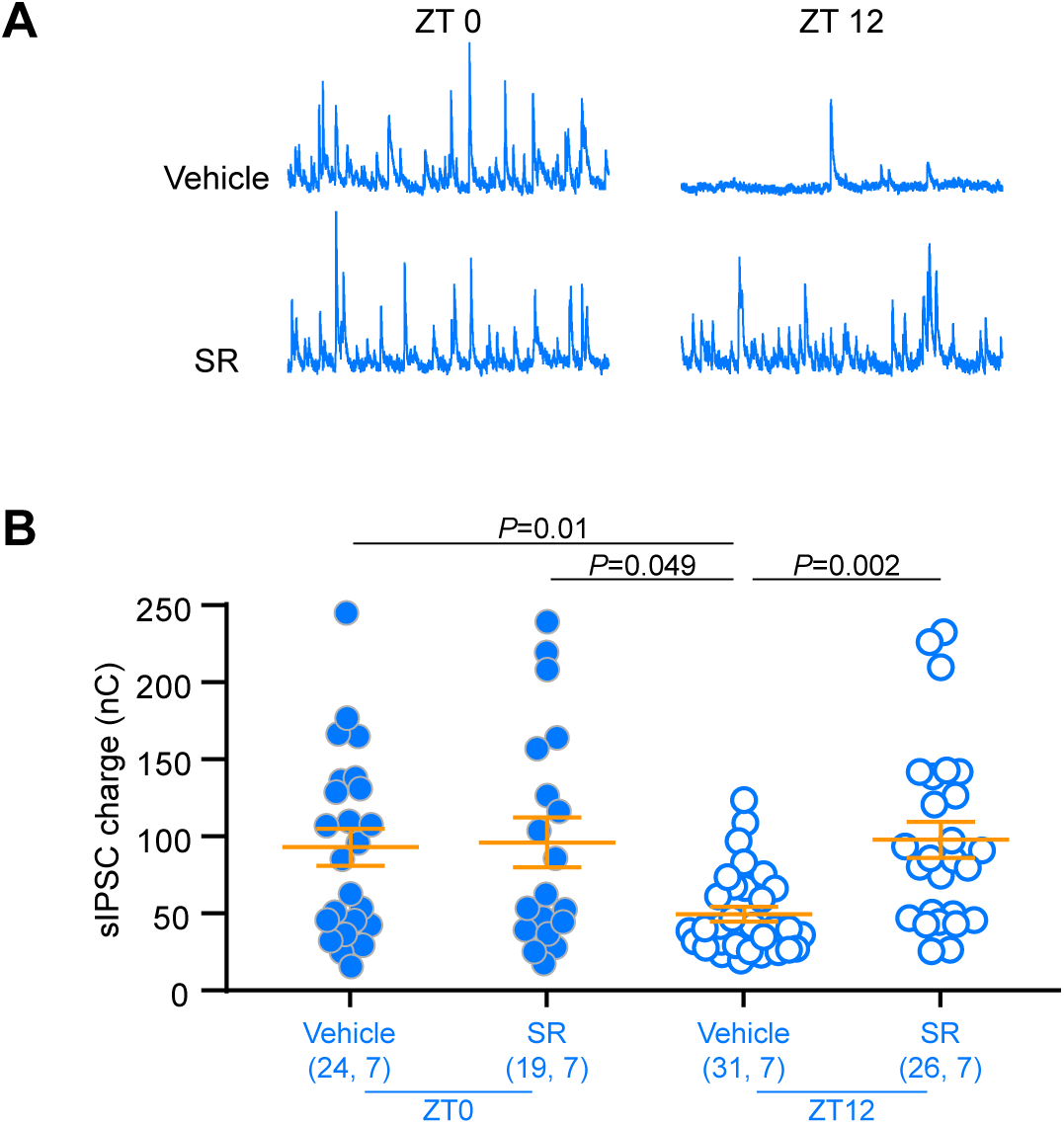
eCB signaling is phase-shifted in BTBR mice. Acute brain slices containing V1 were obtained at ZT0 and ZT12, and sIPSCs in layer 2/3 pyramidal cells were recorded. (A) Example sIPSC traces. (B) Quantification of sIPSCs in the presence and absence of the eCB antagonist SR (10μM). SR enhances inhibitory transmission only during the light phase. Kruskal-Wallis ANOVA on ranks *H*=16.4, *P*=0.001. Dunn’s post-hoc test *P* values are indicated. Sample size is indicated as (cells, mice) and bars represent mean ± SEM.

## Supplementary Tables

**Supplementary Table 1.**
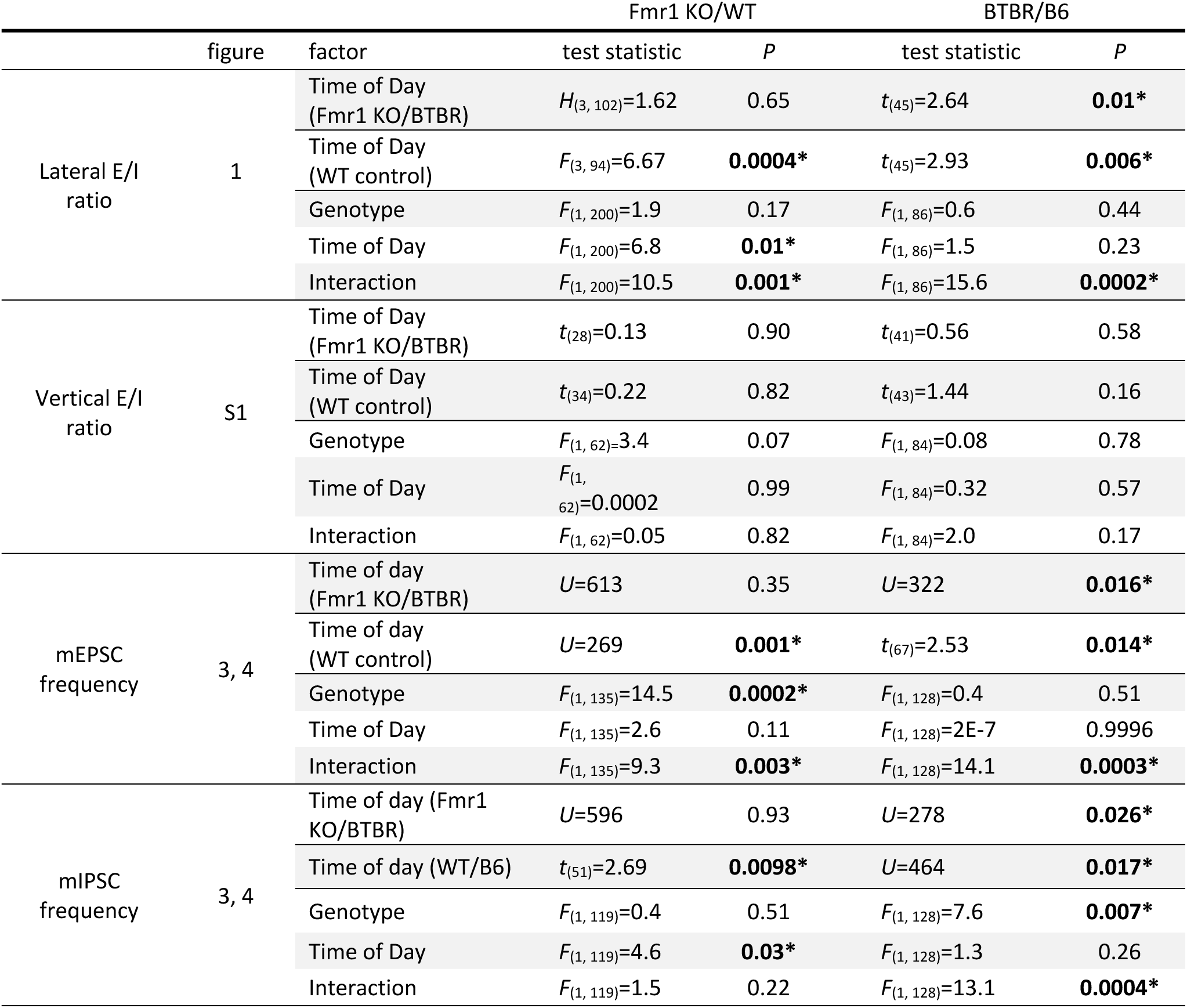
Excitatory/inhibitory synaptic transmission test statistics. Time-of-day changes within each genotype were detected using 1-way ANOVAs and *t* tests (BTBR/B6). For each mouse model, 2-way ANOVAs with genotype (ASD-related vs WT) and time of day as factors were also conducted. *Fmr1* lateral E/I ratio data were pooled within the dark (ZT0, 18) and light (ZT6, 12) phases for the 2-way ANOVA only. Data figures corresponding to each test are indicated. *t*: Student’s *t* test; *U*: Mann-Whitney U test; *H*: Kruskal-Wallis ANOVA on Ranks; *F*: 1-or 2-way ANOVA.

**Supplementary Table 2.**
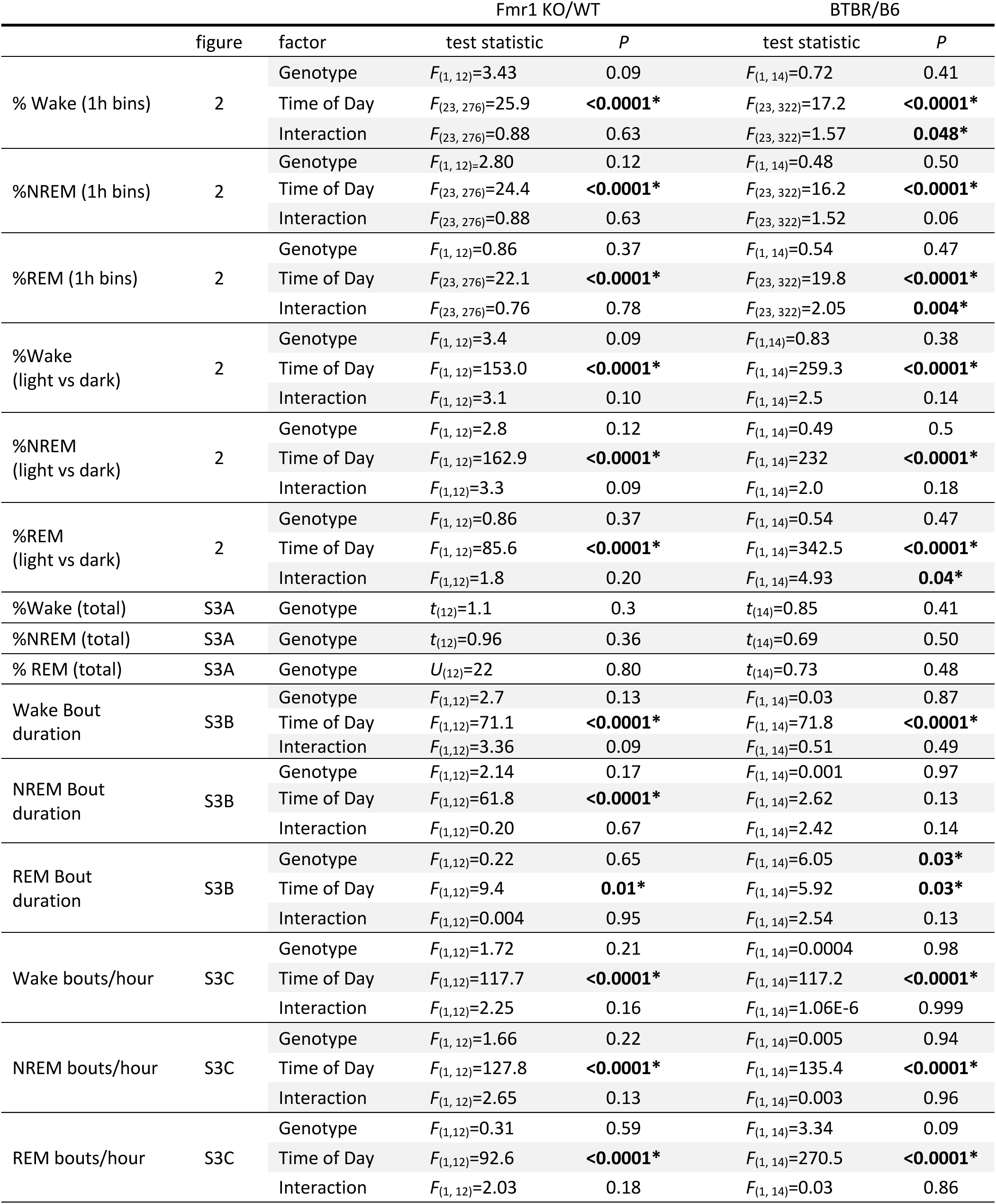
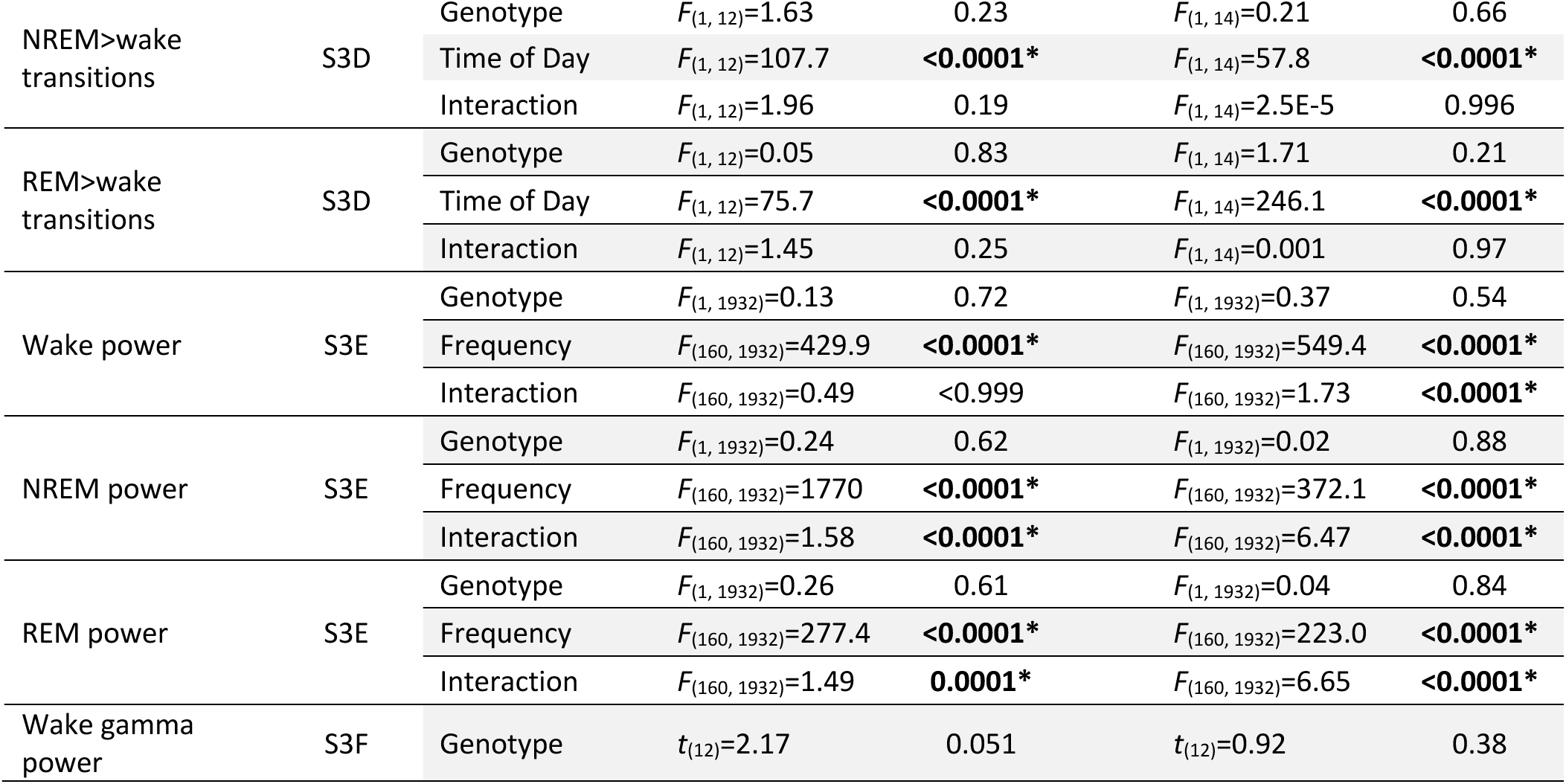
Test statistics for sleep analyses. *F*: 2-way RM ANOVA; *t*: 2-tailed Student’s *t* test, *U*: Mann-Whitney *U* test.

**Supplementary Table 3.**
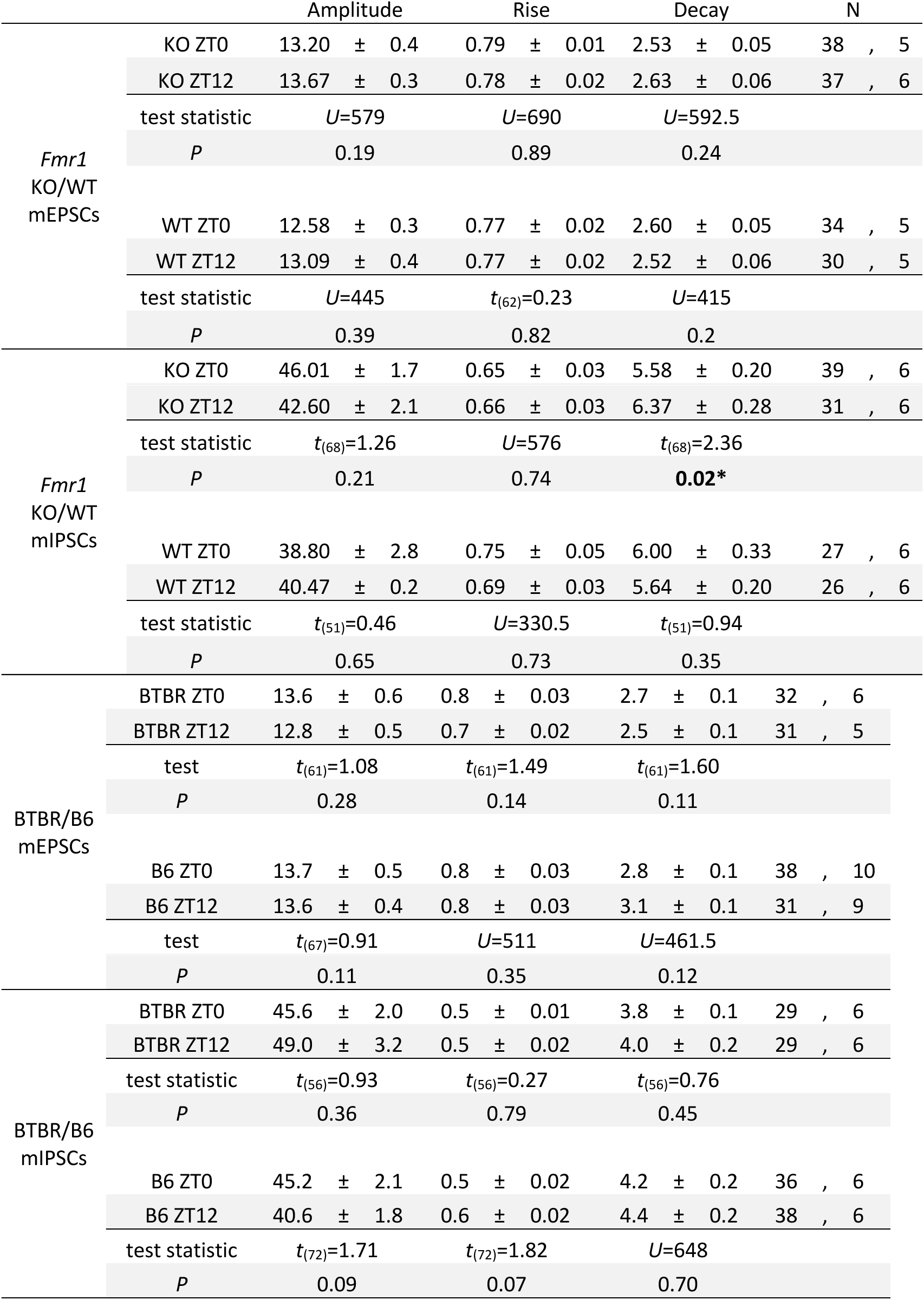
mE/IPSC characteristics (corresponding to Figure 3,4). Groups were compared using *t* or Mann-Whitney *U* tests, as indicated. Data are shown as mean ± SEM.

