## supplementary tables and figures for "Daily oscillation of the excitation/inhibition ratio is disrupted in two mouse models of autism"

**Supplementary Table 1.** Excitatory/inhibitory synaptic transmission test statistics. Time-of-day changes within each genotype were detected using 1-way ANOVAs and *t* tests (BTBR/B6). For each mouse model, 2-way ANOVAs with genotype (ASD-related vs WT) and time of day as factors were also conducted. *Fmr1* lateral E/I ratio data were pooled within the dark (ZT0, 18) and light (ZT6, 12) phases for the 2-way ANOVA only. Data figures corresponding to each test are indicated. *t*: Student's *t* test; *U*: Mann-Whitney U test; *H*: Kruskal-Wallis ANOVA on Ranks; *F*: 1- or 2-way ANOVA.

| | | | Fmr1 KO/WT | | BTBR/B6 | |
|---|---|---|---|---|---|---|
| | figure | factor | test statistic | *P* | test statistic | *P* |
| Lateral E/I ratio | 1 | Time of Day (Fmr1 KO/BTBR) | $H_{(3, 102)}$=1.62 | 0.65 | $t_{(45)}$=2.64 | **0.01*** |
| | | Time of Day (WT control) | $F_{(3, 94)}$=6.67 | **0.0004*** | $t_{(45)}$=2.93 | **0.006*** |
| | | Genotype | $F_{(1, 200)}$=1.9 | 0.17 | $F_{(1, 86)}$=0.6 | 0.44 |
| | | Time of Day | $F_{(1, 200)}$=6.8 | **0.01*** | $F_{(1, 86)}$=1.5 | 0.23 |
| | | Interaction | $F_{(1, 200)}$=10.5 | **0.001*** | $F_{(1, 86)}$=15.6 | **0.0002*** |
| Vertical E/I ratio | S1 | Time of Day (Fmr1 KO/BTBR) | $t_{(28)}$=0.13 | 0.90 | $t_{(41)}$=0.56 | 0.58 |
| | | Time of Day (WT control) | $t_{(34)}$=0.22 | 0.82 | $t_{(43)}$=1.44 | 0.16 |
| | | Genotype | $F_{(1, 62)}$=3.4 | 0.07 | $F_{(1, 84)}$=0.08 | 0.78 |
| | | Time of Day | $F_{(1, 62)}$=0.0002 | 0.99 | $F_{(1, 84)}$=0.32 | 0.57 |
| | | Interaction | $F_{(1, 62)}$=0.05 | 0.82 | $F_{(1, 84)}$=2.0 | 0.17 |
| mEPSC frequency | 3, 4 | Time of day (Fmr1 KO/BTBR) | *U*=613 | 0.35 | *U*=322 | **0.016*** |
| | | Time of day (WT control) | *U*=269 | **0.001*** | $t_{(67)}$=2.53 | **0.014*** |
| | | Genotype | $F_{(1, 135)}$=14.5 | **0.0002*** | $F_{(1, 128)}$=0.4 | 0.51 |
| | | Time of Day | $F_{(1, 135)}$=2.6 | 0.11 | $F_{(1, 128)}$=2E-7 | 0.9996 |
| | | Interaction | $F_{(1, 135)}$=9.3 | **0.003*** | $F_{(1, 128)}$=14.1 | **0.0003*** |
| mIPSC frequency | 3, 4 | Time of day (Fmr1 KO/BTBR) | *U*=596 | 0.93 | *U*=278 | **0.026*** |
| | | Time of day (WT/B6) | $t_{(51)}$=2.69 | **0.0098*** | *U*=464 | **0.017*** |
| | | Genotype | $F_{(1, 119)}$=0.4 | 0.51 | $F_{(1, 128)}$=7.6 | **0.007*** |
| | | Time of Day | $F_{(1, 119)}$=4.6 | **0.03*** | $F_{(1, 128)}$=1.3 | 0.26 |
| | | Interaction | $F_{(1, 119)}$=1.5 | 0.22 | $F_{(1, 128)}$=13.1 | **0.0004*** |

**Supplementary Table 2.** Test statistics for sleep analyses. *F*: 2-way RM ANOVA; *t*: 2-tailed Student's *t* test, *U*: Mann-Whitney *U* test.

| | | | Fmr1 KO/WT | | BTBR/B6 | |
|---|---|---|---|---|---|---|
| | figure | factor | test statistic | *P* | test statistic | *P* |
| % Wake (1h bins) | 2 | Genotype | $F_{(1, 12)}$=3.43 | 0.09 | $F_{(1, 14)}$=0.72 | 0.41 |
| | | Time of Day | $F_{(23, 276)}$=25.9 | **<0.0001*** | $F_{(23, 322)}$=17.2 | **<0.0001*** |
| | | Interaction | $F_{(23, 276)}$=0.88 | 0.63 | $F_{(23, 322)}$=1.57 | **0.048*** |
| %NREM (1h bins) | 2 | Genotype | $F_{(1, 12)=}$2.80 | 0.12 | $F_{(1, 14)}$=0.48 | 0.50 |
| | | Time of Day | $F_{(23, 276)}$=24.4 | **<0.0001*** | $F_{(23, 322)}$=16.2 | **<0.0001*** |
| | | Interaction | $F_{(23, 276)}$=0.88 | 0.63 | $F_{(23, 322)}$=1.52 | 0.06 |
| %REM (1h bins) | 2 | Genotype | $F_{(1, 12)}$=0.86 | 0.37 | $F_{(1, 14)}$=0.54 | 0.47 |
| | | Time of Day | $F_{(23, 276)}$=22.1 | **<0.0001*** | $F_{(23, 322)}$=19.8 | **<0.0001*** |
| | | Interaction | $F_{(23, 276)}$=0.76 | 0.78 | $F_{(23, 322)}$=2.05 | **0.004*** |
| %Wake (light vs dark) | 2 | Genotype | $F_{(1, 12)}$=3.4 | 0.09 | $F_{(1,14)}$=0.83 | 0.38 |
| | | Time of Day | $F_{(1, 12)}$=153.0 | **<0.0001*** | $F_{(1, 14)}$=259.3 | **<0.0001*** |
| | | Interaction | $F_{(1, 12)}$=3.1 | 0.10 | $F_{(1, 14)}$=2.5 | 0.14 |
| %NREM (light vs dark) | 2 | Genotype | $F_{(1, 12)}$=2.8 | 0.12 | $F_{(1, 14)}$=0.49 | 0.5 |
| | | Time of Day | $F_{(1, 12)}$=162.9 | **<0.0001*** | $F_{(1, 14)}$=232 | **<0.0001*** |
| | | Interaction | $F_{(1,12)}$=3.3 | 0.09 | $F_{(1, 14)}$=2.0 | 0.18 |
| %REM (light vs dark) | 2 | Genotype | $F_{(1, 12)}$=0.86 | 0.37 | $F_{(1, 14)}$=0.54 | 0.47 |
| | | Time of Day | $F_{(1, 12)}$=85.6 | **<0.0001*** | $F_{(1, 14)}$=342.5 | **<0.0001*** |
| | | Interaction | $F_{(1,12)}$=1.8 | 0.20 | $F_{(1, 14)}$=4.93 | **0.04*** |
| %Wake (total) | S3A | Genotype | $t_{(12)}$=1.1 | 0.3 | $t_{(14)}$=0.85 | 0.41 |
| %NREM (total) | S3A | Genotype | $t_{(12)}$=0.96 | 0.36 | $t_{(14)}$=0.69 | 0.50 |
| % REM (total) | S3A | Genotype | $U_{(12)}$=22 | 0.80 | $t_{(14)}$=0.73 | 0.48 |
| Wake Bout duration | S3B | Genotype | $F_{(1,12)}$=2.7 | 0.13 | $F_{(1, 14)}$=0.03 | 0.87 |
| | | Time of Day | $F_{(1,12)}$=71.1 | **<0.0001*** | $F_{(1, 14)}$=71.8 | **<0.0001*** |
| | | Interaction | $F_{(1,12)}$=3.36 | 0.09 | $F_{(1, 14)}$=0.51 | 0.49 |
| NREM Bout duration | S3B | Genotype | $F_{(1,12)}$=2.14 | 0.17 | $F_{(1, 14)}$=0.001 | 0.97 |
| | | Time of Day | $F_{(1,12)}$=61.8 | **<0.0001*** | $F_{(1, 14)}$=2.62 | 0.13 |
| | | Interaction | $F_{(1,12)}$=0.20 | 0.67 | $F_{(1, 14)}$=2.42 | 0.14 |
| REM Bout duration | S3B | Genotype | $F_{(1,12)}$=0.22 | 0.65 | $F_{(1, 14)}$=6.05 | **0.03*** |
| | | Time of Day | $F_{(1,12)}$=9.4 | **0.01*** | $F_{(1, 14)}$=5.92 | **0.03*** |
| | | Interaction | $F_{(1,12)}$=0.004 | 0.95 | $F_{(1, 14)}$=2.54 | 0.13 |
| Wake bouts/hour | S3C | Genotype | $F_{(1,12)}$=1.72 | 0.21 | $F_{(1, 14)}$=0.0004 | 0.98 |
| | | Time of Day | $F_{(1,12)}$=117.7 | **<0.0001*** | $F_{(1, 14)}$=117.2 | **<0.0001*** |
| | | Interaction | $F_{(1,12)}$=2.25 | 0.16 | $F_{(1, 14)}$=1.06E-6 | 0.999 |
| NREM bouts/hour | S3C | Genotype | $F_{(1, 12)}$=1.66 | 0.22 | $F_{(1, 14)}$=0.005 | 0.94 |
| | | Time of Day | $F_{(1, 12)}$=127.8 | **<0.0001*** | $F_{(1, 14)}$=135.4 | **<0.0001*** |
| | | Interaction | $F_{(1, 12)}$=2.65 | 0.13 | $F_{(1, 14)}$=0.003 | 0.96 |
| REM bouts/hour | S3C | Genotype | $F_{(1,12)}$=0.31 | 0.59 | $F_{(1, 14)}$=3.34 | 0.09 |
| | | Time of Day | $F_{(1,12)}$=92.6 | **<0.0001*** | $F_{(1, 14)}$=270.5 | **<0.0001*** |
| | | Interaction | $F_{(1, 12)}$=2.03 | 0.18 | $F_{(1, 14)}$=0.03 | 0.86 |

| | | | | | | |
|---|---|---|---|---|---|---|
| NREM>wake transitions | S3D | Genotype | $F_{(1, 12)}$=1.63 | 0.23 | $F_{(1, 14)}$=0.21 | 0.66 |
| | | Time of Day | $F_{(1, 12)}$=107.7 | **<0.0001*** | $F_{(1, 14)}$=57.8 | **<0.0001*** |
| | | Interaction | $F_{(1, 12)}$=1.96 | 0.19 | $F_{(1, 14)}$=2.5E-5 | 0.996 |
| REM>wake transitions | S3D | Genotype | $F_{(1, 12)}$=0.05 | 0.83 | $F_{(1, 14)}$=1.71 | 0.21 |
| | | Time of Day | $F_{(1, 12)}$=75.7 | **<0.0001*** | $F_{(1, 14)}$=246.1 | **<0.0001*** |
| | | Interaction | $F_{(1, 12)}$=1.45 | 0.25 | $F_{(1, 14)}$=0.001 | 0.97 |
| Wake power | S3E | Genotype | $F_{(1, 1932)}$=0.13 | 0.72 | $F_{(1, 1932)}$=0.37 | 0.54 |
| | | Frequency | $F_{(160, 1932)}$=429.9 | **<0.0001*** | $F_{(160, 1932)}$=549.4 | **<0.0001*** |
| | | Interaction | $F_{(160, 1932)}$=0.49 | <0.999 | $F_{(160, 1932)}$=1.73 | **<0.0001*** |
| NREM power | S3E | Genotype | $F_{(1, 1932)}$=0.24 | 0.62 | $F_{(1, 1932)}$=0.02 | 0.88 |
| | | Frequency | $F_{(160, 1932)}$=1770 | **<0.0001*** | $F_{(160, 1932)}$=372.1 | **<0.0001*** |
| | | Interaction | $F_{(160, 1932)}$=1.58 | **<0.0001*** | $F_{(160, 1932)}$=6.47 | **<0.0001*** |
| REM power | S3E | Genotype | $F_{(1, 1932)}$=0.26 | 0.61 | $F_{(1, 1932)}$=0.04 | 0.84 |
| | | Frequency | $F_{(160, 1932)}$=277.4 | **<0.0001*** | $F_{(160, 1932)}$=223.0 | **<0.0001*** |
| | | Interaction | $F_{(160, 1932)}$=1.49 | **0.0001*** | $F_{(160, 1932)}$=6.65 | **<0.0001*** |
| Wake gamma power | S3F | Genotype | $t_{(12)}$=2.17 | 0.051 | $t_{(12)}$=0.92 | 0.38 |

**Supplementary Table 3.** mE/IPSC characteristics (corresponding to Figure 3,4). Groups were compared using *t* or Mann-Whitney *U* tests, as indicated. Data are shown as mean ± SEM.

| | | Amplitude | Rise | Decay | N |
|---|---|---|---|---|---|
| *Fmr1* KO/WT mEPSCs | KO ZT0 | 13.20 ± 0.4 | 0.79 ± 0.01 | 2.53 ± 0.05 | 38 , 5 |
| | KO ZT12 | 13.67 ± 0.3 | 0.78 ± 0.02 | 2.63 ± 0.06 | 37 , 6 |
| | test statistic | $U$=579 | $U$=690 | $U$=592.5 | |
| | *P* | 0.19 | 0.89 | 0.24 | |
| | WT ZT0 | 12.58 ± 0.3 | 0.77 ± 0.02 | 2.60 ± 0.05 | 34 , 5 |
| | WT ZT12 | 13.09 ± 0.4 | 0.77 ± 0.02 | 2.52 ± 0.06 | 30 , 5 |
| | test statistic | $U$=445 | $t_{(62)}$=0.23 | $U$=415 | |
| | *P* | 0.39 | 0.82 | 0.2 | |
| *Fmr1* KO/WT mIPSCs | KO ZT0 | 46.01 ± 1.7 | 0.65 ± 0.03 | 5.58 ± 0.20 | 39 , 6 |
| | KO ZT12 | 42.60 ± 2.1 | 0.66 ± 0.03 | 6.37 ± 0.28 | 31 , 6 |
| | test statistic | $t_{(68)}$=1.26 | $U$=576 | $t_{(68)}$=2.36 | |
| | *P* | 0.21 | 0.74 | **0.02*** | |
| | WT ZT0 | 38.80 ± 2.8 | 0.75 ± 0.05 | 6.00 ± 0.33 | 27 , 6 |
| | WT ZT12 | 40.47 ± 0.2 | 0.69 ± 0.03 | 5.64 ± 0.20 | 26 , 6 |
| | test statistic | $t_{(51)}$=0.46 | $U$=330.5 | $t_{(51)}$=0.94 | |
| | *P* | 0.65 | 0.73 | 0.35 | |
| BTBR/B6 mEPSCs | BTBR ZT0 | 13.6 ± 0.6 | 0.8 ± 0.03 | 2.7 ± 0.1 | 32 , 6 |
| | BTBR ZT12 | 12.8 ± 0.5 | 0.7 ± 0.02 | 2.5 ± 0.1 | 31 , 5 |
| | test | $t_{(61)}$=1.08 | $t_{(61)}$=1.49 | $t_{(61)}$=1.60 | |
| | *P* | 0.28 | 0.14 | 0.11 | |
| | B6 ZT0 | 13.7 ± 0.5 | 0.8 ± 0.03 | 2.8 ± 0.1 | 38 , 10 |
| | B6 ZT12 | 13.6 ± 0.4 | 0.8 ± 0.03 | 3.1 ± 0.1 | 31 , 9 |
| | test | $t_{(67)}$=0.91 | $U$=511 | $U$=461.5 | |
| | *P* | 0.11 | 0.35 | 0.12 | |
| BTBR/B6 mIPSCs | BTBR ZT0 | 45.6 ± 2.0 | 0.5 ± 0.01 | 3.8 ± 0.1 | 29 , 6 |
| | BTBR ZT12 | 49.0 ± 3.2 | 0.5 ± 0.02 | 4.0 ± 0.2 | 29 , 6 |
| | test statistic | $t_{(56)}$=0.93 | $t_{(56)}$=0.27 | $t_{(56)}$=0.76 | |
| | *P* | 0.36 | 0.79 | 0.45 | |
| | B6 ZT0 | 45.2 ± 2.1 | 0.5 ± 0.02 | 4.2 ± 0.2 | 36 , 6 |
| | B6 ZT12 | 40.6 ± 1.8 | 0.6 ± 0.02 | 4.4 ± 0.2 | 38 , 6 |
| | test statistic | $t_{(72)}$=1.71 | $t_{(72)}$=1.82 | $U$=648 | |
| | *P* | 0.09 | 0.07 | 0.70 | |

Supplementary Fig 1.

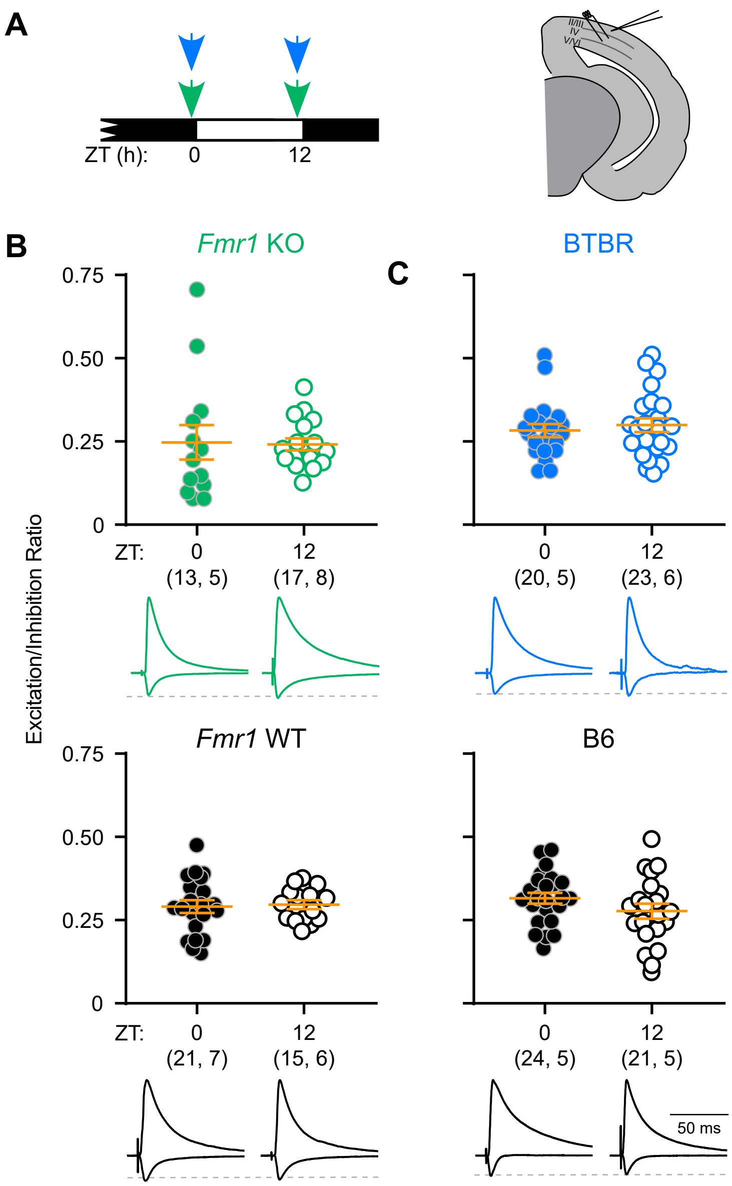

Supplementary Figure 2.

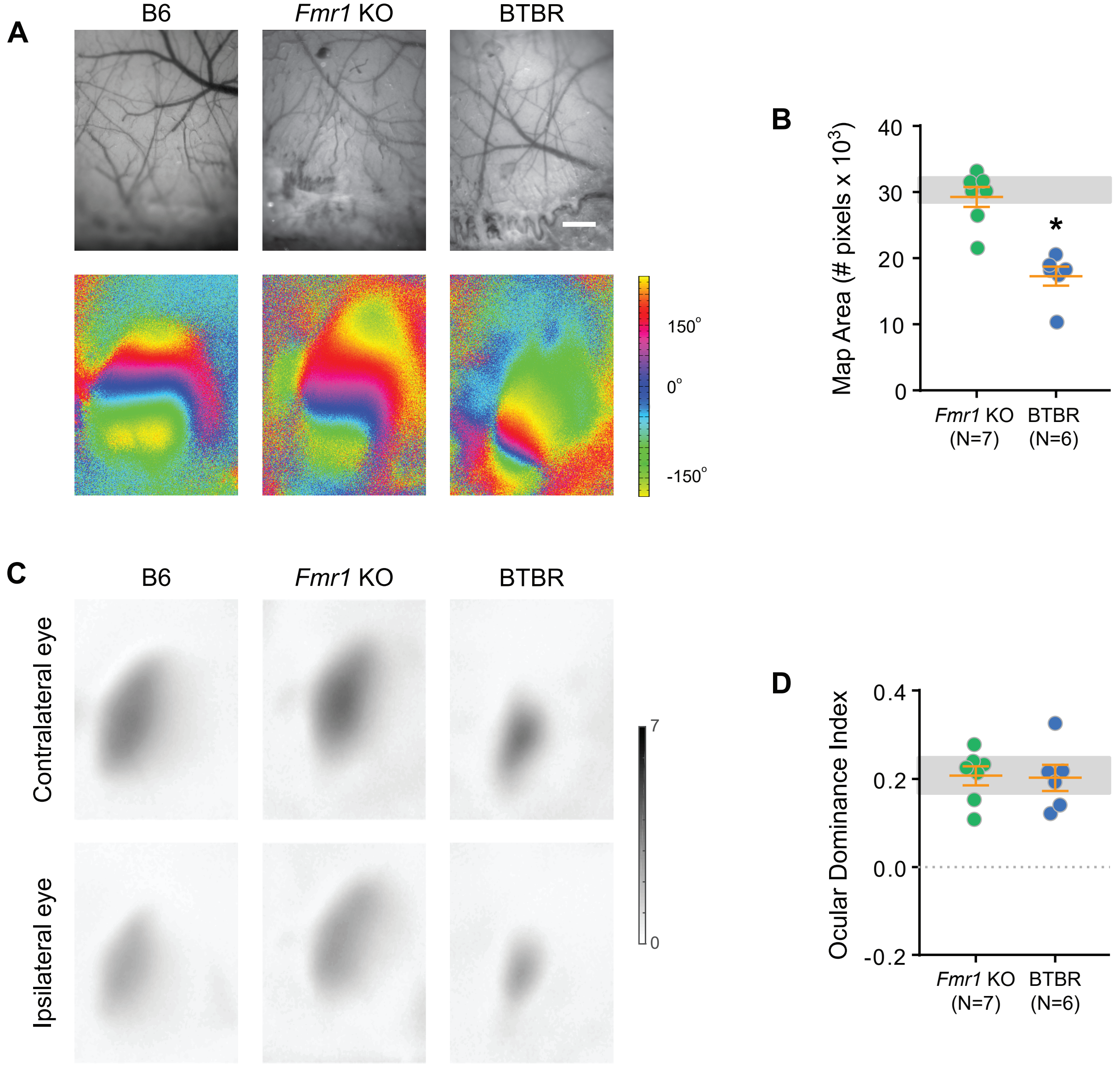

Supplementary Figure 3.

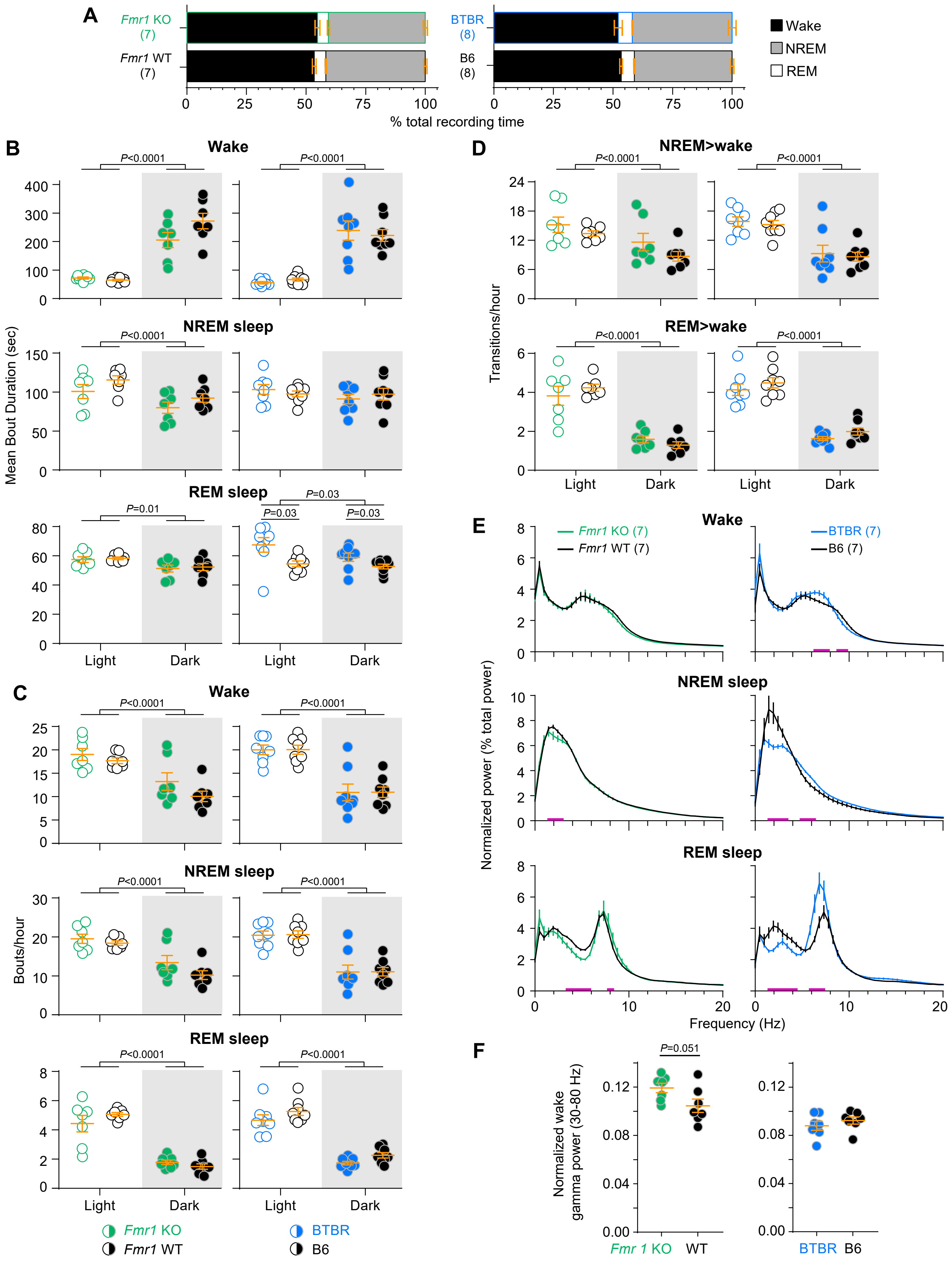

Supplementary Figure 4.

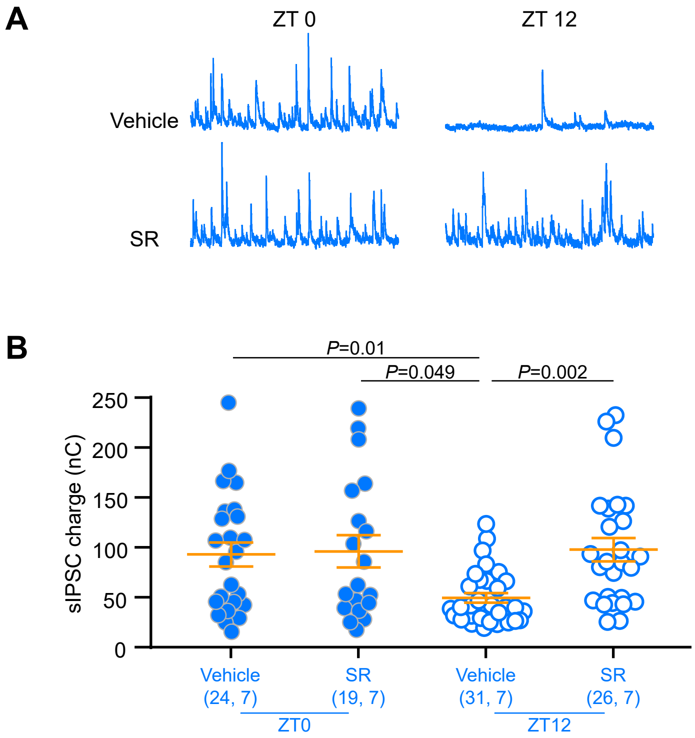
